# Upregulated GIRK2 counteracts ethanol-induced changes in excitability & respiration in human neurons

**DOI:** 10.1101/2023.03.22.533236

**Authors:** Iya Prytkova, Yiyuan Liu, Michael Fernando, Isabel Gameiro-Ros, Dina Popova, Chella Kamarajan, Xiaoling Xuei, David B. Chorlian, Howard J. Edenberg, Jay A. Tischfield, Bernice Porjesz, Zhiping P. Pang, Ronald P. Hart, Alison Goate, Paul A. Slesinger

**Author notes:** Corresponding authors: Contact information: Paul Slesinger, Alison Goate.

## Abstract

Genome-wide association analysis (GWAS) of electroencephalographic endophenotypes for alcohol use disorder (AUD) has identified non-coding polymorphisms within the *KCNJ6* gene. *KCNJ6* encodes GIRK2, a subunit of a G protein-coupled inwardly-rectifying potassium channel that regulates neuronal excitability. How changes in GIRK2 affect human neuronal excitability and the response to repeated ethanol exposure is poorly understood. Here, we studied the effect of upregulating *KCNJ6* using an isogenic approach with human glutamatergic neurons derived from induced pluripotent stem cells (male and female donors). Using multi-electrode-arrays, population calcium imaging, single-cell patch-clamp electrophysiology, and mitochondrial stress tests, we find that elevated GIRK2 acts in concert with 7-21 days of ethanol exposure to inhibit neuronal activity, to counteract ethanol-induced increases in glutamate response, and to promote an increase intrinsic excitability. Furthermore, elevated GIRK2 prevented ethanol-dependent changes in basal and activity-dependent mitochondrial respiration. These data support a role for GIRK2 in mitigating the effects of ethanol and a previously unknown connection to mitochondrial function in human glutamatergic neurons.

**SIGNIFICANCE STATEMENT:** Alcohol use disorder (AUD) is a major health problem that has worsened since COVID, affecting over 100 million people worldwide. While it is known that heritability contributes to AUD, specific genes and their role in neuronal function remain poorly understood, especially in humans. In the current manuscript, we focused on the inwardly-rectifying potassium channel GIRK2, which has been identified in an AUD-endophenotype genome-wide association study. We used human excitatory neurons derived from healthy donors to study the impact of GIRK2 expression. Our results reveal that elevated GIRK2 counteracts ethanol-induced increases in glutamate response and intracellular calcium, as well as deficits in activity-dependent mitochondrial respiration. The role of GIRK2 in mitigating ethanol-induced hyper-glutamatergic and mitochondrial offers therapeutic promise for treating AUD.

## INTRODUCTION

A genome wide association study (GWAS) conducted by the Collaborative Study on the Genetics of Alcoholism (COGA) identified single nucleotide polymorphisms (SNPs) in the *KCNJ6* gene associated with variations of electroencephalogram (EEG) frontal theta event-related oscillations (θ-EROs) to task-relevant target stimuli during a visual oddball task, in addition to other measures of attention, inhibitory control, and reward processing (Kang et al., 2012; Kamarajan et al., 2017). Low frontal θ-ERO during such tasks is characteristic of individuals with alcohol use disorder (AUD) as well as their high-risk offspring and considered to be an endophenotype for AUD (Jones et al., 2006; Rangaswamy et al., 2007). *KCNJ6* SNPs have also been linked to frontal θ-ERO developmental trajectories during adolescence and young adulthood (Chorlian et al., 2017). These SNPs are non-coding, except for a single imputed synonymous SNP, and therefore are expected to primarily impact gene expression. A recent study investigated a haplotype of 22 linked, noncoding SNPs in the *KCNJ6* gene and found that they are associated with altered expression of *KCNJ6* transcripts and protein in human neurons (Popova et al., 2023).

*KCNJ6* encodes a G protein-coupled inwardly-rectifying potassium channel subunit 2, GIRK2 (Kir3.2), which plays a key role in controlling neuronal excitability (Lüscher and Slesinger, 2010). In human brains, GIRK2 is largely present as a homotetramer or in a heterotetramer with GIRK1 (Kir3.1) (Lüscher and Slesinger, 2010). The channel is broadly expressed throughout the brain in multiple neuronal types, including cortical pyramidal neurons (Victoria et al., 2016). GIRK2 is coupled to G_i/o_ G-protein coupled receptors (GPCRs), including γ-aminobutyric acid receptor GABA_B_ and Group II metabotropic glutamate receptors (Lüscher and Slesinger, 2010). Ligand-binding and subsequent dissociation of the G-protein trimer leads to GIRK activation by Gβγ, outward potassium flux, and inhibition of neuronal firing (Lüscher and Slesinger, 2010; Glaaser and Slesinger, 2015). Importantly, alcohol can directly activate GIRK2 via a hydrophobic pocket in the cytoplasmic domain (Aryal et al., 2009; Bodhinathan and Slesinger, 2014; Glaaser and Slesinger, 2015). In addition to these direct mechanisms, GIRK channels depend on phosphatidylinositol 4,5-biphosphate (PIP_2_)-binding and membrane cholesterol to facilitate conformational changes underlying channel activation (Glaaser and Slesinger, 2015, 2017). Studies in mice have shown that GIRK2 affects reward mechanisms and behaviors associated with alcohol and other drugs of abuse (Blednov et al., 2001a, 2001b; Hill et al., 2003).

The direct impact of altered GIRK2 expression on human neurons remains poorly understood. Conceivably, different GIRK2 levels can contribute to changes in neuronal excitability, influencing θ-EROs. To better understand the effect of GIRK2 levels independent of *KCNJ6* SNP haplotypes and AUD genetics, we manipulated the expression of GIRK2 either through CRISPRa (Ho et al., 2017) or with a lentivirus in human glutamatergic neurons derived from healthy donors, facilitating a comparison within the same cell line (i.e., isogenic). We focused on glutamatergic neurons because of the well-documented effects of ethanol on glutamate signaling (Läck et al., 2007; Kim et al., 2010; Das et al., 2016), the frontal cortex associations of GWAS findings (Kang et al., 2012), and SNP expression trait quantitative trait loci effects (https://www.gtexportal.org/home/snp/rs2835872). We hypothesized that elevated GIRK2 expression would influence neuronal adaptations to ethanol exposure.

We used bulk RNAseq to characterize early response to ethanol in glutamatergic neurons with endogenous GIRK2 levels, following a 7-day intermittent ethanol exposure (IEE) protocol (Scarnati et al., 2020). We observed downregulation of genes involved in neuronal development and enrichment of mitochondrial and metabolic pathways. We then tracked the developmental effects of extended (7-21 days) IEE combined with increased GIRK2 expression (↑GIRK2), using multi-electrode arrays (MEA), calcium imaging, patch-clamp electrophysiology, and mitochondrial stress tests to assess glutamatergic function and activity-dependent energy demands. We followed-up with bulk RNAseq of 21-day IEE iso-CTL and ↑GIRK2 neurons to identify potential molecular mechanisms underlying the differential adaptations in glutamate sensitivity, membrane excitability, and cellular respiration. Together, our results begin to elucidate the relationship between GIRK2, ethanol, glutamatergic signaling, and mitochondrial health.

## MATERIALS AND METHODS

### RESOURCE AVAILABILITY

#### Lead Contact

Further information and requests for resources and reagents should be directed to and will be fulfilled by the lead contact Paul A. Slesinger.

#### Materials availability

This study did not generate new unique reagents.

#### Data and code availability

- RNAseq data were deposited at NCBI GEO and are publicly available as of date of publication. Accession numbers: GSE226746; GSE244985.
- Any additional information required to reanalyze data reported in this paper is available from the lead contact upon request.

### EXPERIMENTAL MODEL DETAILS

#### Astrocytes

Primary human fetal astrocytes (HFAs) were obtained from a commercial vendor and cultured in astrocyte medium according to the manufacturer specifications (ScienCell, Cat#: 1801). After cells have reached their terminal duplication capacity, they were transplanted onto 12mm glass acid-etched coverslips coated in 80ug/mL Matrigel (Thermo Fisher Scientific, Cat#: 08-774-552) and cultured for a week to achieve appropriate coverage and attachment.

#### Neural progenitor cells (NPCs) and induced neurons (iNs)

NPCs and dCas9-VPR NPCs from de-identified control hiPSC cell lines used in this study were derived from both female (cell-lines 12455 and 9429) and male (cell-lines BJ, 3440, 553-VPR, 2607-VPR) donors (Topol et al., 2016; Ho et al., 2017; Tcw et al., 2017). NPCs and dCas9-NPCs were cultured in DMEM/F12 with sodium pyruvate GlutaMAX medium (Thermo Fisher Scientific, Cat#: 10565018) supplemented with FGF2 (Bio-Techne, Cat#: 233-FB), B27 (Life Technologies, Cat#: 17504-044), N2(Life Technologies, Cat#: 17502-048), and penicillin-streptomycin (Thermo Fisher Scientific, Cat#: 15-140-122). NPCs were induced into neurons with doxycycline-inducible transcription factor *NGN2* with neomycin antibiotic selection (Addgene #99378), following the protocol described by (Ho et al., 2016). Antibiotic selection was performed on Day 1 post-induction with 10 μg/mL neomycin sulfate (BioVision Incorporated, Cat#: 9620). Starting at Day 2 post-induction iNs were cultured in Neurobasal medium (Life Technologies, Cat#: 21103-049) supplemented with N2 and penicillin-streptomycin. At day 7 post-induction, iNs were infected with *KCNJ6* vector or *KCNJ6* gRNAs in dCas9-VPR expressing cell lines to generate ↑GIRK2 neurons. CRISPRa iso-CTLs were generated using a scramble guide RNA. Lenti cohort controls did not receive any additional viral vector. At Day 10, iNs were dissociated using the Papain Dissociation System following the manufacturer’s protocol (Worthington Biochemical Cat#: LK003153) and then transplanted onto HFA-covered coverslips with thiazovivin (1μg/mL) (EMD-Millipore, Cat#: 420220) to facilitate attachment. Starting at Day 14, B27 Plus culture system medium (Thermo Fisher Scientific, Cat#: A36534-01) supplemented with 2% fetal bovine serum (FBS) (Sigma Aldrich, Cat#: F3135) and 1% penicillin/streptomycin (Thermo Fisher Scientific, Cat#: 15-140-122) was introduced through half medium changes every other day to promote electrophysiological maturation.

### METHOD DETAILS

#### KCNJ6 gRNA Design and Cloning

Single gRNAs were designed using CRISPR-ERA (http://crispr-era.stanford.edu/), with the predicted top six ranked guides selected for in-vitro validation. For lentiviral cloning, IDT synthesized oligonucleotides were annealed and phosphorylated using PNK (NEB#C3019) at 37° C for 30min, 95° C for 5min and ramp-down to 25° C at 5C/min. Annealed oligos were cloned into lentiGuide-TdTomato-Hygro (Addgene#99376) using BsmB1-v1 (NEB, discontinued) in an all-in-one T7 ligase (NEB#M0318S) based Golden-Gate assembly (37° C for 5min, 20°C for 5min x15 cycles), and subsequently transformed into NEB10beta (NEB#C3019). Purified DNA from propagated colonies were confirmed for gRNA insertion via Sanger sequencing (Genewiz). Following Sanger sequencing confirmation, gRNAs were prepped for lenti-virus production.

#### Lentivirus preparation

Lentiviral vectors were prepared using PEI transfection of HEK-293T cells as previously described (Tiscornia et al., 2006). Viral titer was concentrated to 10^8^ IU/mL. Vectors used are listed in Key Resources Table (Addgene #99378, #19780, #99347, #99376, #197032).

#### Intermittent Ethanol Exposure

Starting at Day 21 post-induction, 100% ethanol (EtOH) (Sigma Aldrich, Cat# e7023) was added to B27 Plus System culture medium to achieve a final concentration of 17 mM. A previous study demonstrated that after 24 h, ethanol concentration in medium is reduced to ∼5 mM, therefore medium was “spiked” with 1ul EtOH/ 1.5 mL medium daily to maintain ∼17 mM concentration (Halikere et al., 2020; Scarnati et al., 2020).

#### Bulk RNAseq

*NGN2-*induced neurons were scraped from 12-well plates at 28 or 42 days of differentiation, spun down, and frozen. Total RNA extraction from frozen cell pellets, library preparation, and sequencing was performed by Genewiz (Azenta US, Inc., Indianapolis, IN). Total RNA was extracted from fresh frozen cell pellet samples using Qiagen RNeasy Plus Universal mini kit following manufacturer’s instructions (Qiagen, Hilden, Germany). RNA samples were quantified using Qubit 2.0 Fluorometer (Life Technologies, Carlsbad, CA, USA) and RNA integrity was checked using Agilent TapeStation 4200 (Agilent Technologies, Palo Alto, CA, USA). RNA sequencing libraries were prepared using the NEB-Next Ultra RNA Library Prep Kit for Illumina using manufacturer’s instructions (NEB, Ipswich, MA, USA). The samples were sequenced on the Illumina 4000 instrument, using a 2×150bp Paired End (PE) configuration. Image analysis and base calling were conducted by the Control software. Raw sequence data (.bcl files) generated the sequencer were converted into fastq files and de-multiplexed using Illumina’s bcl2fastq 2.17 software.

Raw reads were assessed for quality using FastQC (version 0.11.8) before and after adapter sequence trimming by Cutadapt (version 4.1). Reads were pseudo-aligned using Salmon (version 0.13.1) for estimation of normalized gene expression (TPM, i.e., transcript per million). Reads were also aligned to the human reference genome (Gencode version 28) and quantified for gene read counts, both in STAR (version 2.5.3a). In day-28 neurons’ differential gene expression analyses, read counts were modelled in the linear mixed model using batch, sex and donor information as the random effect terms and ethanol-treatment/control as the fixed effect term by the R (version 4.1.0) package “variancePartition” (version 1.28.0). In Day 42 neurons, differential gene expression analysis was conducted using the NOISeq package in R (version 2.42.0) (Tarazona et al., 2015), correcting for batch and donor effects, with ↑GIRK2 + IEE groupwise comparison to untreated isogenic controls. In day-28 neurons, gene set enrichment analysis (GSEA) was performed with 1,000 permutations using R package GOtest (github.com/mw201608/Gotest) in the human molecular signatures database ‘C2.CP’,’C5.BP’,’C5.CC’ and ‘C5.MF’ collections. In day-42 neurons synaptic pathway analysis was performed using SynGO online portal (version 1.2) (Koopmans et al., 2019).

#### Western blot

Tissue from 28-day-old neurons was collected by scraping and membrane protein was extracted using MemPer ThermoFisher kit (Thermo Fisher Scientific, Cat#: 89842) according to the manufacturer’s instructions. The membrane fraction was then denatured for 20 minutes at 70°C in MES running buffer (Life Technologies, Cat#: B0002) and separated on a BOLT^TM^ 4-12% Bis-Tris 1mm gel (Thermo Fisher Scientific, Invitrogen, Cat#: NW04125BOX) at 200mV for 20-30 minutes. Transfer to a nitrocellulose membrane was achieved using iBlot2. Membranes were then blocked in 5% milk in PBST for 1 hour. To ensure antibody specificity, antigen-competition was utilized as a control, where the antibody (Alomone, Cat #: APC-006) was pre-incubated with the antigen in the blocking solution for one hour on ice; 1:3 antibody to antigen ratio was used. Anti-NaK ATPase antibody (Cell Signaling Technology, Cat#: 23565S) was used as a loading control. Membranes were incubated in 1:500 antibody or antibody/antigen mixture overnight at 4°C. Anti-rabbit-HRP secondary antibody (Cell Signaling Technology, Cat#: 7074P2) at 1:1000 dilution was incubated for 1 hour at room temperature. Western-bright peroxidase (Advansta, Cat#: K-12045-D20) 2-minute incubation and 90s exposure on UVP imager were used to obtain images.

#### Immunocytochemistry

Neurons at day 28 were fixed in methanol for twenty minutes at −20°C, followed by permeabilization in 0.2% Triton-X in PBS (Mg^2+^, Ca^2+^), and blocking in 2% bovine serum albumin, 5% normal goat serum, 2% Triton-X in PBS (Mg^2+^, Ca^2+^) at room temperature. Primary antibodies were diluted in 4% NGS, 0.1% Triton in PBS (no Mg^2+^, Ca^2+^) (GIRK2 (Alomone, Cat #: APC-006) 1:400, TUJ1 (Biolegend, Cat#: 801213, RRID: AB_2728521) 1:1000) and incubated overnight at 4°C. Secondary antibodies were incubated for two hours at room temperature at 1:500 dilution [Anti-Rabbit AlexaFluor-488 (Thermo Fisher Scientific, Cat#: A-11008, RRID: AB_143165), Anti-Mouse AlexaFluor-647 (Thermo Fisher Scientific, Cat#: A-21235, RRID: AB_2535804)]. Images were acquired at the Icahn School of Medicine Microscopy and Advanced Biomaging Core, using the Zeiss Upright LSM-780, Axio Imager 2 confocal microscope with the 100x objective. Maximum intensity projection was obtained from Z-stacks consisting of 5-15 6.6 μm slices, using Zeiss Zen Black Software. The maximum intensity projection image was analyzed using Analyze Particles in FIJI (ImageJ) (Schindelin et al., 2012).

#### Multi Electrode Array (MEA)

CytoView 48-well MEA plates (Axion Biosystems, M768-tMEA-48B) were coated with 80ug/mL Matrigel (Thermo Fisher Scientific, Cat#: 08-774-552) and seeded with human fetal astrocytes (ScienCell, Cat#: 1801). Neurons were transplanted on Day 10 of induction. The plate was loaded onto the Axion Maestro MEA reader (Icahn School of Medicine at Mount Sinai Stem Cell Engineering Core), and the electrical activity was analyzed using AxIS 2.0 Neuronal Module Software. Ten minute MEA recordings (Ho et al., 2016; Soni et al., 2021) were acquired on Days 28, 35, and 42 post-induction, at approximately 24 h after last ethanol supplementation (see **Figure 3B**), which is estimated to be ∼ 5 mM after ∼24h (Scarnati et al (2020)). Wells with little or no activity (< 0.01 Hz) were excluded from analysis. One outlier data point was identified by the Grubbs outlier test and removed from further analysis (**Figure 3D**).

#### Patch clamp electrophysiology

Day 42 neurons were recorded at room-temperature (∼20°C) in artificial cerebrospinal fluid (ACSF) external solution consisting of 125 mM NaCl, 5 mM KCl, 10 mM D-Glucose, 10 mM HEPES-Na, 3.1 mM CaCl_2,_ and 1.3 mM MgCl_2_6·H_2_O. Solution pH was adjusted to 7.4 with NaOH and filtered with 0.22 µM PEI bottle-top filter. Osmolarity was measured at 290-300 mOsm with VAPRO® Vapor Pressure Osmometer (Fisher Scientific, ELITechGroup Model 5600, Cat#: NC0044806). Patch pipettes were pulled to a resistance of 4-4.5 MΩ. GIRK current was recorded with an internal solution containing 140 mM K-D-Gluconate, 4 mM NaCl, 2 mM MgCl_2_6·H_2_O, 1.1 mM EGTA, 5 mM HEPES, 2 mM Na_2_ATP, 5 mM Na-Creatine-PO_4_, and 100 μM GTPyS. pH adjusted to 7.4, and osmolarity was measured at 300-310 mOsm. EPSCs were recorded gap-free at −70 mV using voltage-clamp with an internal solution of 135 mM CsCl2, mM 10 HEPES-Na, 1 mM EGTA, 1 Na mM–GTP, and 1 mM QX-314 as described by Yang et al., 2017 (Yang et al., 2017). All electrophysiology chemicals purchased from Sigma.

Resting membrane potential was obtained in current-clamp mode (I = 0). Cells with a resting membrane potential higher than −40 mV and a membrane resistance lower than 100 MΩ were excluded from further recordings and analyses. For measuring excitability, the membrane potential was adjusted to −55 mV (equivalent to −72 mV with a junction-potential correction of 17 mV). Neuronal excitability was measured with current injection steps of 20 pA (up to 16 steps). Rheobase was defined as the voltage at which the first current injection step elicited an action potential first. Neurons with fewer than two action potential spikes were excluded from further analysis. In voltage-clamp, GIRK currents were assessed using a 100 ms voltage ramp (from −100 to 0 mV) to examine inward rectification of GIRK channels, followed by 300 ms pulse at −40 mV to measure the outward GIRK current, and a 20 ms step to −45 mV to monitor membrane resistance, from a holding potential of −40 mV. Voltage pulses were delivered every 2 s at a sampling rate of 5 kHz, and lowpass filter frequency of 2 kHz. After 4-5 minutes 30 µM SCH-23390 (Fisher Scientific Cat#: 092550) in ACSF was applied for 1 minute, followed by a 2-minute ACSF washout. All data were acquired using the Clampex software (v11.0.3.3, Molecular Devices) with a Axopatch B200 amplifier and Digidata 1550B digitizer with HumSilencer enabled to minimize noise. EPSCs were analyzed with Easy Electrophysiology software (v2.5.2, Easy Electrophysiology). GIRK current quantification and I-clamp data were analyzed in Clampfit software (v10.7.0.3, Molecular Devices). The amplitude of GIRK currents were measured as an average of 20 ms at – 40 mV, approximately 150 ms after the voltage ramp.

#### Seahorse Assay

Assays were conducted at the Icahn School of Medicine Mitochondrial Analysis Facility, using an XFe96 Agilent Seahorse Analyzer. NPCs were plated directly in 96-well Agilent Seahorse Cell Culture plates (Agilent Technologies, Cat#: 101085-004), coated with Matrigel, and induced into neurons the following day. Each cell line was plated in 8-replicate wells for each experiment. On the day of the experiment, neurons were incubated for 1 hour in CO_2_-free incubator in Agilent Seahorse DMEM assay medium (Agilent Technologies, cat# 103575-100) supplemented with 1 mM sodium pyruvate (Agilent Technologies, Cat#: 103578-100), 10 mM glucose (Agilent Technologies, Cat#: 103577-100), and 2 mM glutamine (Agilent Technologies, Cat#:103579-100). Acute 10 μM glutamate injection was applied to four replicate wells/donor/experiment during the mitochondrial stress test (Agilent Technologies, Cat#: 103015-100). The mitochondrial stress test was performed according to the manufacturer’s protocol, with 1 μM oligomycin, 1 μM FCCP, 1 μM rotenone/antimycin-A. Data were acquired in Seahorse Wave software (Agilent Technologies).

After conclusion of the experiment, medium was aspirated and cells were fixed with Methylene blue overnight at 4° C. The dye was rinsed out the following morning with distilled water and cells were lysed with 4% acetic acid and 40% methanol. The lysate was transferred to a clear flat-bottom 96-well plate to measure absorbance at 595 nm using a Varioskan plate reader (Thermo Scientific, Cat#: VL0000D0) and SkanIt Software for Microplate Readers, ver. 6.0.2.3 (Thermo Scientific, Cat#: 5187139). Absorbance values were applied as a scaling factor for raw oxygen consumption rate (pmol/min) to normalize respiratory rate to cell density. Normalized raw data were exported as an Excel spreadsheet for further analysis in R Studio, v3.6.1 (R Core Team, 2020). Mitochondrial respiration parameters were calculated according to the mitochondrial stress test assay user manual. Each experiment replicate is reported as the average of 4 wells.

#### Calcium Imaging

Neurons were incubated in 4 µM Fluo4-AM in ACSF (same solution as for patch clamp electrophysiology) for 20 minutes at 37° C, protected from light. Cells were then destained for an additional 5 minutes in ACSF at room temperature prior to imaging. Fluorescence was measured with at 10x objective on a Nikon TE2000 inverted microscope, 480 nm LED, excitation filter of (485 nm) and emission filter (530 nm). Images were obtained with a sCMOS Zyla 5.5 camera (Oxford Instruments, Andor), using a 10-fps rate and 4×4 binning under constant perfusion of ACSF in a laminar flow diamond-shaped chamber (Model #RC-25; Warner Instruments) at RT (∼20° C). Spontaneous activity was recorded for 3 minutes. 10 μM glutamate in ACSF was applied in 30 second pulses, with a 1-minute ACSF washout period. 15 mM KCl in ACSF was applied at the end of the recording for 1 minute, followed by a 1-minute ACSF washout. Time of stimulus application was tagged during the recording.

Raw fluorescence data were collected using Nikon Elements software (NIS Elements AR; version 5.20.01). A background region of interest (ROI) was identified to subtract noise. ROIs were identified using automatic detection in Nikon Elements. All ROIs were manually curated and raw fluorescence data were exported into Excel. Subsequent analysis steps were carried out in R Studio (v3.6.1 of R). Each fluorescence trace was corrected for baseline-drift using a penalized least-squares algorithm with lambda = 10 (AirPLS) (Zhang et al., 2010) and ΔF/F_0_ was calculated according to the formula (F_t_-F_0_)/F_0_, where F_0_ was defined as the minimum fluorescence intensity (RFU) in the first 10s of the recording. Traces were filtered with a 3^rd^ order Butterworth filter using “signal” package version 0.7-6 (Ligges et al., 2021) and spikes were detected using “Pracma” package version 2.3.3 (Borchers, 2022) (**Figure 4-1).** A neuronal spike was defined as <30s in duration and above 5 standard deviations of the background ROI. ROIs with astrocyte-like spikes (lasting > 30s) were excluded from the analyses. ROIs without any detectable spikes were also excluded.

### EXPERIMENTAL DESIGN AND STATISTICAL ANALYSES

Sample size of 4-6 donors and isogenic design was selected to maximize sensitivity and minimize the false discovery rate, as shown by Germain and Testa, 2017. The experimenter was not blinded to experimental conditions during data collection and analysis. All statistical analyses were carried out in R Studio, v3.6.1 (R Core Team, 2020), using the following packages for quantitation and visualization: jtools version 2.1.4 (Long, 2022), lme4 version 1.1-27.1 (Bates et al., 2022), ggplot2 version 3.3.5 (Wickham et al., 2023). One-tailed student’s t-test was used for immunocytochemistry puncta % μm^2^ quantitation, with a p = 0.05 significance cutoff. Generalized Linear Mixed Models (GLMM) were used for patch-clamp electrophysiology, MEA, calcium imaging, and mitochondrial stress test data to account for donor and batch replicate effects. GLMM analysis maintains appropriate biological grouping of variables (i.e., donors) and can account for random variance between replicate experiments, while still retaining the full power of multiple measurements (e.g., neuronal ROIs in calcium imaging) (see (Popova et al., 2023)). Additionally, utilizing GLMM permits the specification of different distributions of data, unlike ANOVA, which relies on the assumption of a normal distribution (see (Yu et al., 2022)). Data normality was tested using the Shapiro-Wilk test. Random effects included donor cell line, neuronal differentiation batch, and/or experimental replicates (e.g., coverslips for calcium imaging and patch-clamp electrophysiology data). Fixed variables included GIRK2 expression level and ethanol exposure, tested for separate and interaction effects, with a p = 0.05 significance cutoff. Model fit was selected based on lowest AIC (Akaike information criterion) values. Poisson distribution was specified for all spike count data. Representative formula: Spikes ∼ GIRK2*EtOH + (1|Donor) + (1|Batch), family = “poisson”. All bar and line plots include +/- standard error of the mean, with individual data-points shown for each replicate and color-coded by donor.

## RESULTS

### Enhancing GIRK2 currents in human neurons using CRISPRa and lentivirus expression of KCNJ6

To study the effect of increased GIRK2 expression on neurogenin 2 (*NGN2*)-induced glutamatergic neurons (Ho et al., 2016 p.2) and their response to chronic ethanol exposure, we employed two different methodologies: dCas9-VPR activation (CRISPRa (Ho et al., 2017)) of the *KCNJ6* gene and expression from a lentivirus. CRISPRa has the advantage of upregulating the endogenously-encoded *KCNJ6* gene, but it relies on stable expression of the dCas9-coupled transcriptional activator VPR and chromatin accessibility of the endogenous gene (Ho et al., 2017; Savell et al., 2018). Lentiviral vector-mediated expression bypasses these limitations, but expresses the *KCNJ6* mRNA without the 3’UTR, which may play a key role in protein trafficking to the plasma membrane as well as transcript stability (Loya et al., 2008).

We therefore evaluated the effect of both strategies on GIRK2 expression and function in *NGN2*-induced neurons using qPCR, Western blotting, and immunocytochemistry. The endogenous levels of GIRK2 protein appeared low, with average puncta density of 2.2 ± 0.5% of neuronal marker β-III tubulin (TUJ1)-immunoreactive area (μm^2^). CRISPRa was accomplished using cell line 553 stably expressing dCas9-VPR (Ho et al., 2017) and co-expression of three pooled gRNAs targeting the *KCNJ6* promoter (see Methods) (**Figure 1A-B**). In comparison to 553/dCas9-VPR alone, GIRK2 puncta density increased to 23 ± 3% (p = 0.009) (N = 9 iso-CTL, 16 ↑GIRK2) of TUJ1 area in 553/dCas9-VPR + *KCNJ6* gRNAs (CRISPRa). For lentiviral (Lenti) expression of *KCNJ6*, we used a lentivirus containing the *KCNJ6* gene with mCherry red fluorescent marker in cell line 12455 (control line from Knight Alzheimer’s Disease Research Center) (Tcw et al., 2017). Here, we observed an increase in GIRK2 puncta density to 24 ± 6% (p = 0.005) (N = 9 iso-CTL, 10 ↑GIRK2) of TUJ1 area (**Figure 1D-F**). Thus, both CRISPRa- and Lenti-mediated *KCNJ6* expression appeared to increase the levels of GIRK2 protein expression similarly (**Figure 1C-F**).

**Figure 1.**
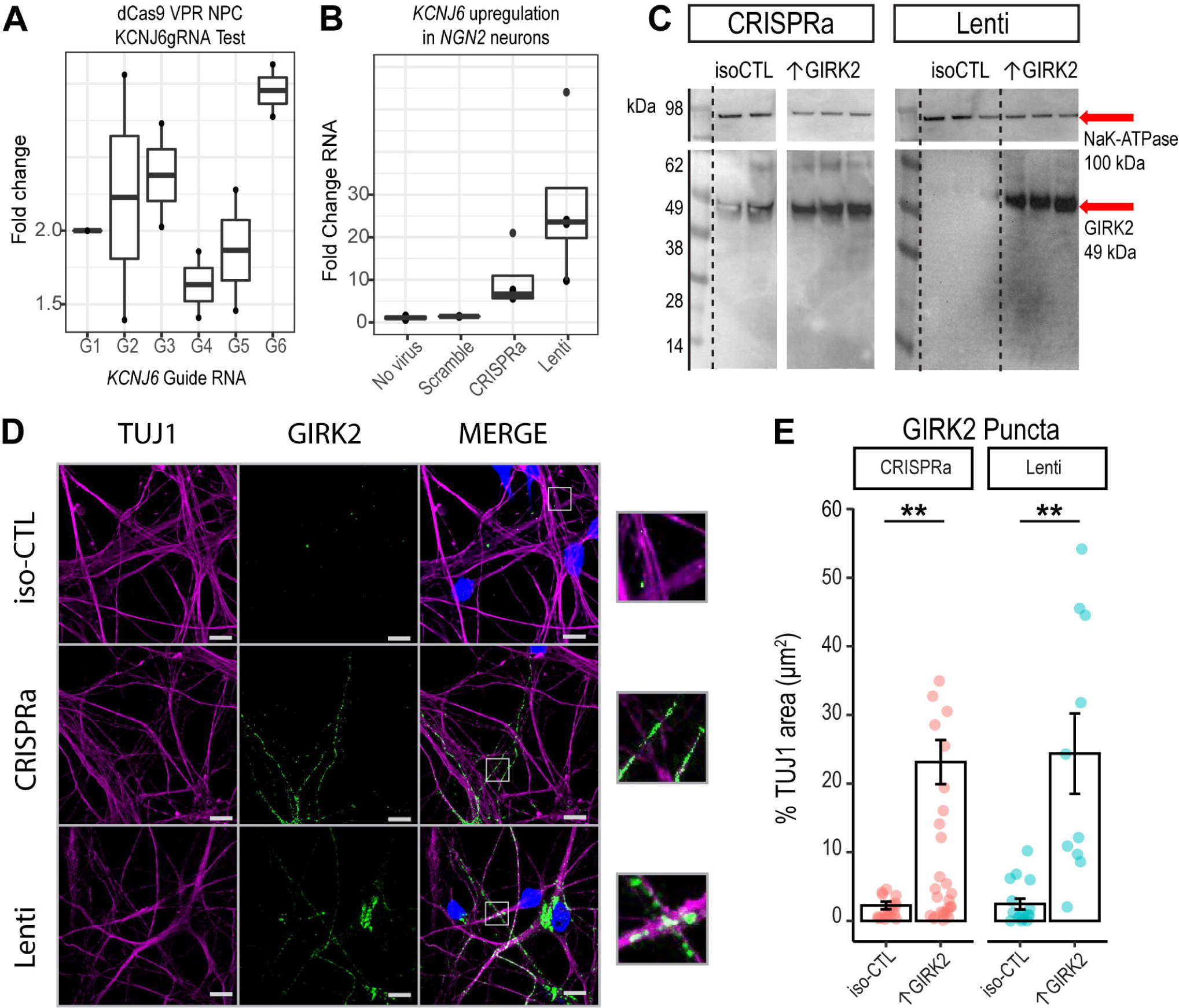
CRISPRa and lentivirus-mediated *KCNJ6* expression similarly increase GIRK2 protein in human glutamatergic neurons. **A.** RT-qPCR test of 6 *KCNJ6*gRNAs normalized to a scramble gRNA control in D42 *NGN2* neurons. (N = 1 donor, 2 replicates). **B.** RT-qPCR of *KCNJ6* expression in D42 NGN2 neurons with scramble gRNA, no virus control, combined *KCNJ6*gRNAs 1, 3, & 6 (CRISPRa) and lentiviral expression (Lenti). (N = 1 donor, 3 replicates)**. C.** Western blot of iso-CTL, CRISPRa, and Lenti expression of GIRK2 (MW = 49kDa, red arrow). NaK-ATPase loading control (MW = 198k Da, red arrow). (N = 1 donor, 2 replicates scr-gRNA control, 3 replicates *KCNJ6* gRNAs, no virus control, and lentivirus *KCNJ6* vector). **D.** Representative images of isogenic control (iso-CTL) neurons, CRISPRa- and Lenti-expression ↑GIRK2 neurons stained for TUJ1 (magenta), DAPI (blue), and GIRK2 (green); 100x magnification; scale bar and insets = 10 μm. **E.** Normalization of GIRK2 puncta μm^2^ to TUJ1 μm^2^; each point represents a single image; coral = CRISPRa and teal = Lenti (N images = CRISPRa: 9 iso-CTL (scr-gRNA control), 16 ↑GIRK2 (*KCNJ6* gRNA); *KCNJ6* lentivirus: 9 iso-CTL (no virus control), 10 ↑GIRK2). All experiments were carried out in Day 42 *NGN2* neurons.

We next examined whether higher protein expression led to an increase in GIRK current, using whole-cell patch-clamp electrophysiology. Both 553/dCas9-VPR and 12455/Lenti neurons were grown on glass coverslips for 5-6 weeks with human fetal astrocytes to improve maturation (Johnson et al., 2007; Tang et al., 2013; Odawara et al., 2014). To study GIRK currents, we included 100 μM GTPyS in the internal solution to activate endogenous G proteins and produce maximal GIRK activation, as previously described (Schreibmayer et al., 1996; Slesinger et al., 1997; Federici et al., 2009). Given that the expression of voltage-gated potassium as well as non-selective ion channels in neurons (Duménieu et al., 2017) can obscure the inward rectification of GIRK channels, we used a GIRK-specific blocker SCH-23390 (referred to as SCH) to assess GIRK current (Kuzhikandathil and Oxford, 2002; Zhao et al., 2020). At ∼5 min following establishment of the whole-cell recording, application of 30 μM SCH rapidly and reversibly inhibited the outward current (**Figure 2A**). The reversal potential for the SCH-inhibited current was approximately −80 mV (**Figure 2B**), which is near the calculated equilibrium potential for potassium (*E*_K_) ([K^+^]_out_ = 5 mM, [K^+^]_in_ = 140 mM), confirming the potassium selectivity of the channel.

**Figure 2.**
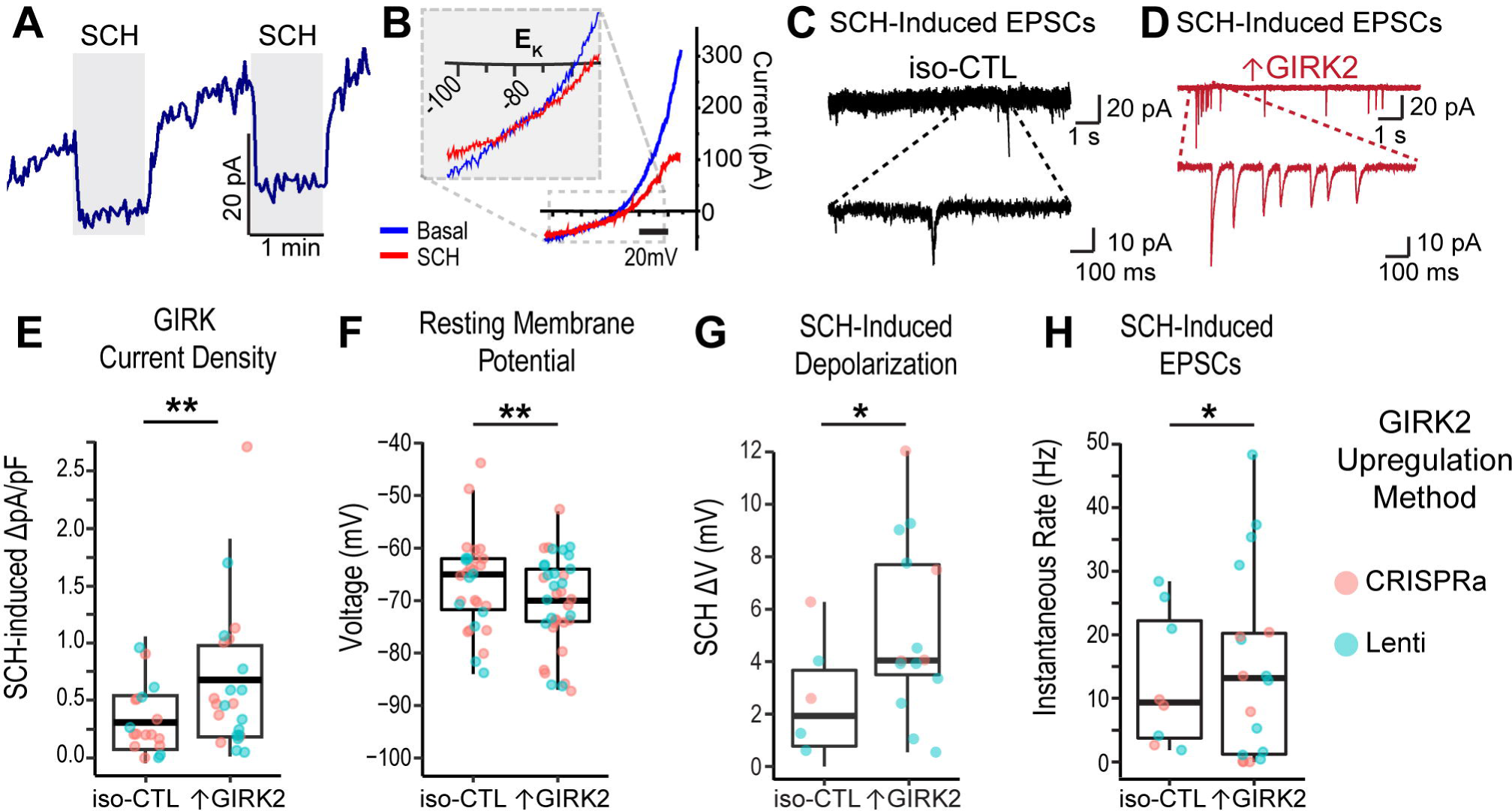
Increased GIRK2 protein expression alters electrophysiology of human glutamatergic neurons. **A.** Representative traces of GTPγS-activated current with two 1-min pulses of 30 μM SCH-23390 (grey rectangles) voltage-clamped at −40 mV. **B.** Representative current-voltage curves of basal (blue) and GIRK2-inhibited (SCH-23390 red trace) currents; inset: magnification of current reversal at the potassium equilibrium potential (E_K_). **C-D:** Representative sEPSCs for iso-CTL (black) (**E**) and ↑GIRK2 (red) (**F**) neurons with SCH pulse. Main trace = 10 s, magnified inset = 1 s **E-H.** Boxplots showiing mean (middle bar), upper and lower quartiles (box), and 1.5 inter-quartile range (whiskers) of (**E**) SCH-inhibited GIRK current density (pA/pF), (N = 19 iso-CTL cells (12 scr-gRNA + 7 no virus); 23 ↑GIRK2 (11 CRISPRa + 12 Lenti)); (**F**) resting membrane potential (N = 29 iso-CTL cells (20 scr-gRNA + 9 no virus); 37↑GIRK2 (20 CRISPRa + 17Lenti)); (**G**) SCH-induced shift in resting membrane potential (N = 6 iso-CTL cells (2 scr-gRNA + 4 no virus); 13 ↑GIRK2 (3 CRISPRa + 10 Lenti)); and (**H**) instantaneous sEPSC rate (Hz) with SCH (N = 8 iso-CTL cells (3 scr-gRNA + 5 no virus); 18↑GIRK2 (7 CRISPRa + 11 Lenti)). Each point represents a cell; point color indicates GIRK2 expression method (coral = CRISPRa; teal = Lenti). Statistics: generalized linear mixed model (GLMM) with expression method and batch random effects. *p < 0.05, ** p < 0.01. See Extended Data Table 2-1 for additional patch properties (membrane capacitance, access resistance, rheobase, and action potential threshold).

Overall, CRISPRa-mediated enhancement and lenti expression of *KCNJ6* led to similar increases in GIRK2 protein (**Figure 1C-F**); we therefore pooled these data and refer to them as ↑GIRK2. The GIRK current density for SCH-inhibited current was significantly larger in ↑GIRK2 neurons, as compared to isogenic controls (+ 0.34 pA/pF; p = 0.028; N = 19 iso-CTL, 23 ↑GIRK2) (**Figure 2E**). Furthermore, ↑GIRK2 neurons exhibited more hyperpolarized resting membrane potentials (–6.7 mV; p = 0.009; N = 29 iso-CTL, 37 ↑GIRK2) (**Figure 1H**), a larger SCH-induced membrane depolarization (+ 3.1 mV; p = 0.016; N = 6 iso-CTL, 13 ↑GIRK2) (**Figure 1I**), and a higher frequency of SCH-induced spontaneous EPSC (sEPSCs), (+ 7.9 Hz; p = 0.021; N = 8 iso-CTL, 18 ↑GIRK2) (**Figure 1E, F, & J**). ↑GIRK2 neurons did not significantly differ in rheobase, suggesting little impact of GIRK2 on action potential thresholds (see **Table 2-1**). Additional neuronal properties, including cell capacitance, membrane resistance, or evoked spiking activity revealed no significant differences (**Table 2-1**), indicating preservation of basic cell properties with ↑GIRK2. Collectively, these data support the conclusion that an increase in GIRK channel expression results in a functional increase in GIRK channel activity.

### 7-day 17 mM ethanol protocol: transcriptomic evidence of downregulated neuronal maturation and upregulated metabolic activity

Prior to investigating how ↑GIRK2 interacts with ethanol to influence neuronal activity, we first sought to investigate the effect of 17 mM ethanol (equivalent to 0.08% blood alcohol concentration) in iso-CTL neurons over 7 days on gene expression as determined by bulk RNAseq. We used an intermittent-ethanol-exposure (IEE) paradigm, described previously (Scarnati et al., 2020; Popova et al., 2023). For IEE, we supplemented ethanol daily to maintain 17 mM concentration, since the ethanol concentration in the cell culture medium decreases to 5 mM or less after 24h (Lieberman et al., 2012; Scarnati et al., 2020; Popova et al., 2023). Glutamatergic neurons from four donors were induced in two batches in the absence of astrocytes to provide a pure neuronal population for RNAseq analyses. At day 21 post induction, when neurons begin to be electrically active (Ho et al., 2016), we treated half of the cultures with the IEE protocol for 7 days (on D21), and harvested the neurons at day 28 post-induction (**Figure 3A**).

**Figure 3.**
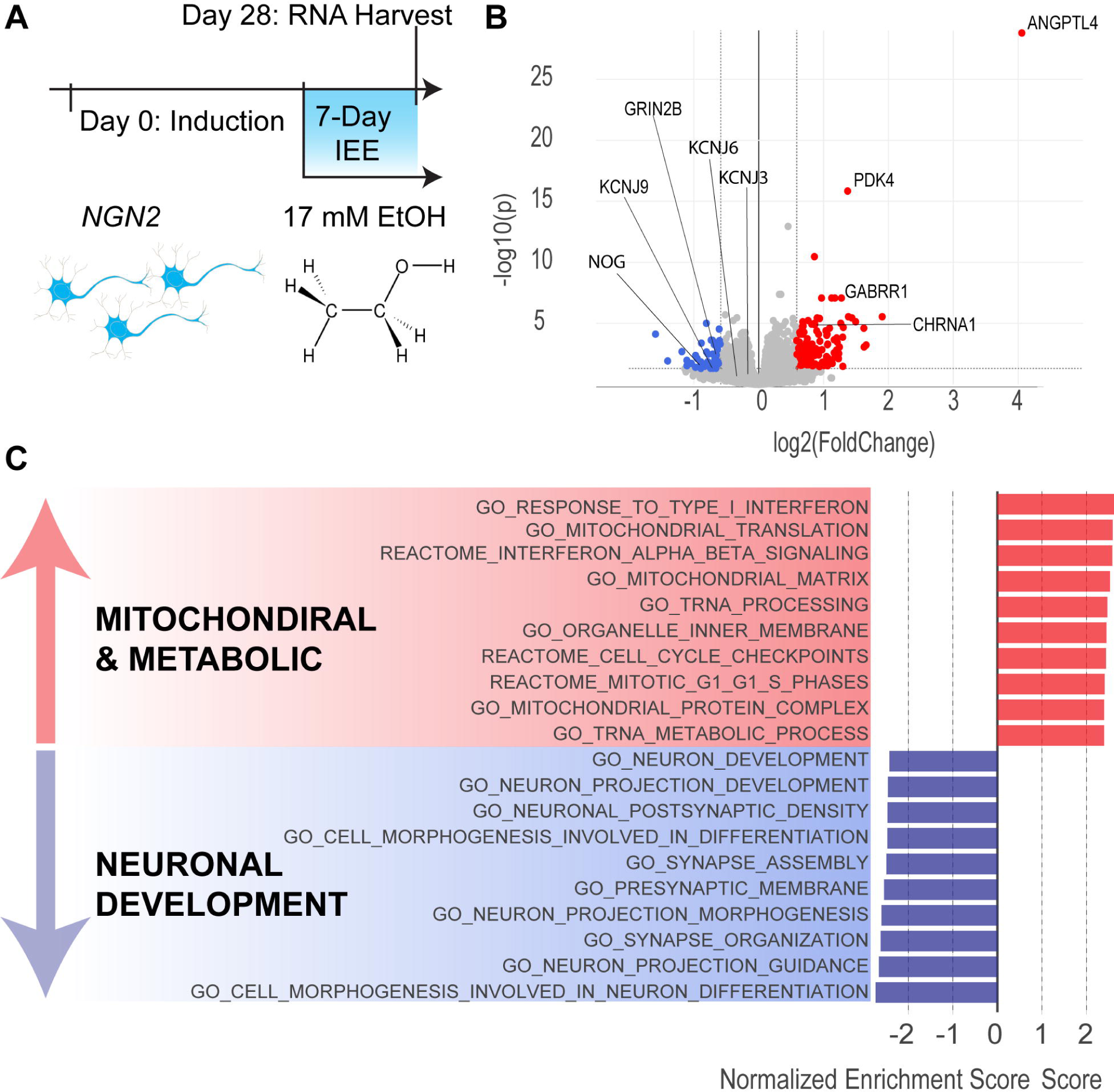
7-Day Intermittent ethanol exposure (IEE) increases expression of mitochondrial and metabolic genes and impairs neuronal development genes. **A.** Timeline for neuronal induction, intermittent 17 mM ethanol exposure, and RNA harvest. **B.** Volcano plot of differentially expressed genes (DEGs) (fold-change = 1.5 and p-value = 0.05 cutoffs). Blue dots represent downregulated genes and red dots represent upregulated genes. **C.** Gene ontology categories of differentially expressed genes. Bars depict normalized enrichment score for each category. See Extended Data Table 3-1 for complete list of significant DEGs. N = 4 donors, 2 batches, 4 replicate wells.

Differential gene expression analysis revealed that 7-day IEE resulted in 127 upregulated and 50 downregulated genes, with Benjamini-Hochberg adjusted p-value cutoff of 0.05 and fold-change cutoff of 1.5 (**Figure 3B**, **Table 3-1**). Among them, *PDK4* and *ANGPTL4* showed the largest increase in fold-change. These two genes are involved in regulating the switch between glycolysis and fatty acid oxidation(La Paglia et al., 2017; Ma et al., 2019) and are both influenced by upstream proliferator-activated receptor-γ (PPARγ) (Cippitelli et al., 2017; La Paglia et al., 2017). Interestingly, *PDK4* overexpression is associated with loss of metabolic flexibility and promotes excessive transport of calcium into mitochondria (Zhang et al., 2014; Ma et al., 2019; Jeon et al., 2020). Gene ontology (GO) analysis shows the robust upregulation of the associated biological processes (**Figure 3D**). Most enriched categories are associated with metabolic, mitochondrial, and inflammatory pathways, suggesting one of the impacts of 7-day IEE is on energy utilization in glutamatergic neurons.

Furthermore, GO analysis revealed a significant depletion of differentially expressed genes involved in neuronal development, morphogenesis, and synapse organization - including neurodevelopmental gene *NOG*, voltage gated potassium channel subunit *KCNH1,* inwardly rectifying potassium channel subunit *KCNJ9*, and ionotropic glutamate receptor subunit *GRIN2B* (**Figure 2D**, **Table 2-1**). Expression of other inwardly rectifying potassium channels, including *KCNJ6* (GIRK2) and *KCNJ3* (GIRK1), and other canonical markers of glutamatergic identity, such as glutamate transporter *SCLA17A7* (vGLUT1) and glutamate receptor *GRIA4* (AMPAR), was not significantly affected. Interestingly, several genes involved in inhibitory neurotransmission were upregulated, including *CHRNA9* and *GABRR1* (**Figure 2C**). Taken together, these data show that 7-day IEE promotes significant alterations in the transcriptome of glutamatergic neurons, indicating the presence of metabolic stress and perturbations in glutamatergic neuronal maturation.

### GIRK2 expression counteracts hyperactivity induced by prolonged (7-21 days) 17 mM ethanol exposure

We hypothesized that ↑GIRK2 and IEE would act in concert to inhibit neuronal activity, because of ethanol-induced downregulation of neuronal maturation genes and the direct activation of GIRK channels (Bodhinathan and Slesinger, 2014). To address this, we first used Multi-Electrode Arrays (MEA) to measure neuronal local field potentials over weeks of ethanol exposure. Using 553/dCas9-VPR and 12455/Lenti neurons, we followed the same 17 mM IEE paradigm used in bulk RNAseq, starting the protocol at day 21, two weeks after the start of ↑GIRK2 expression (**Figure 4A**). To track the effects of ethanol exposure and GIRK2 expression levels on the developmental time-course, we extended IEE duration beyond 7 days and collected MEA data at days 28, 35, and 42 (corresponding with 1, 2 and 3 weeks, respectively, of IEE). To maximize consistency of measurements across the multi-week experiment, MEA recordings were collected at the same time of day, ∼24 h after the last supplementation of ethanol, at a time when ethanol was ∼5 mM in the culture medium (Scarnati et al., 2020) (**Figure 4B**).

**Figure 4.**
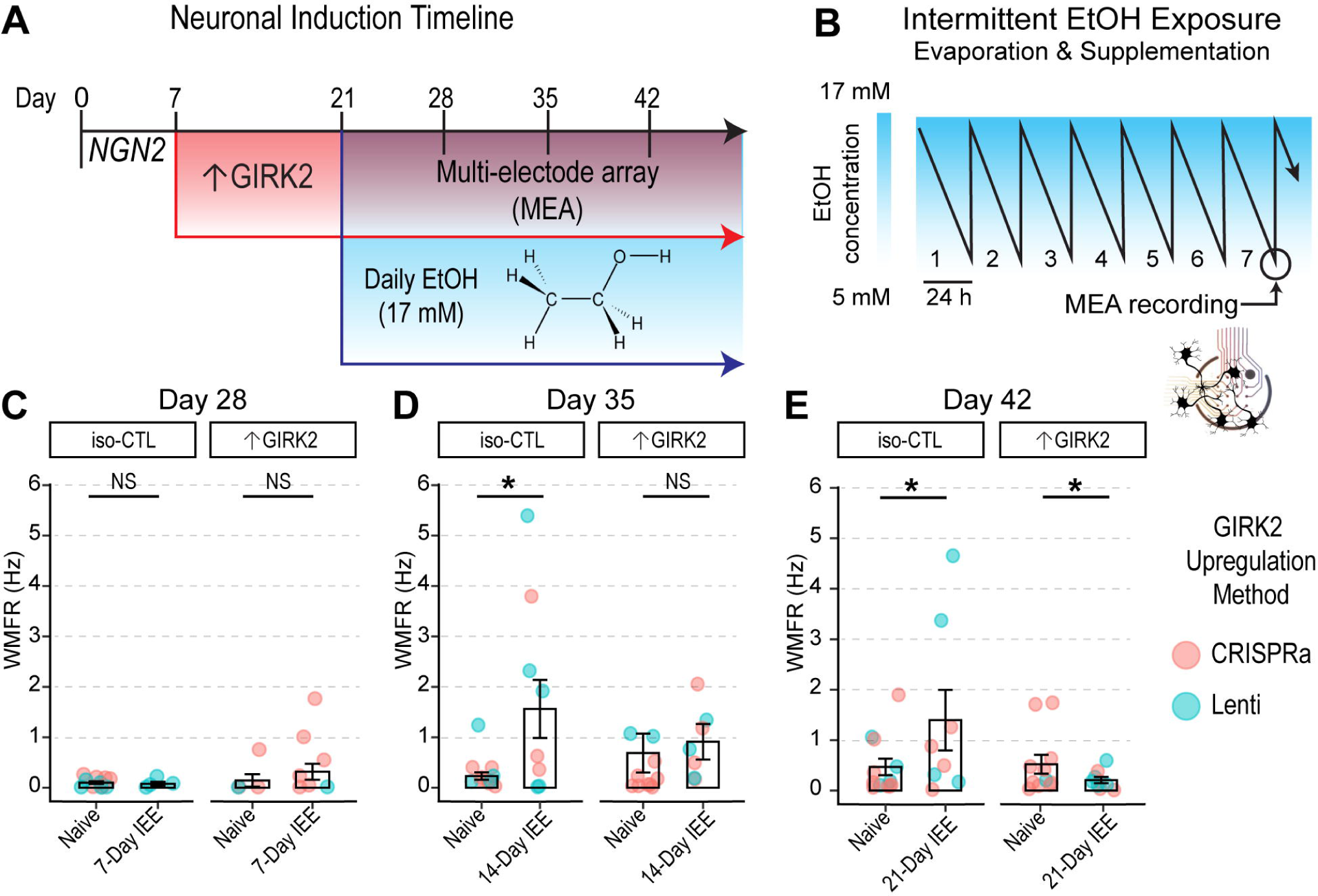
Increased GIRK2 expression counteracts hyperactivity induced by low-ethanol conditions. **A.** Timeline of neuronal differentiation, GIRK2 upregulation and ethanol exposure. **B.** Schematic of 7-day IEE, showing change in ethanol concentration over 24 hours due to evaporation (Scarnati et al., 2020; Popova et al., 2023). **C-E**: Neuronal firing rates (WMFR) measured with multi-electrode array (MEA) is shown for *NGN2* neurons at Day 28 (1 week ethanol**) (C)**, 35 (2 weeks ethanol) **(D)**, and 42 (3 weeks ethanol) **(E)**; each point represents an MEA well; N = 2 donors | 5 batches. Point color indicates GIRK2 expression method (coral = CRISPRa; teal = Lenti). GLMM includes batch and donor random effects. * p < 0.05.

At day 28, MEA recordings showed very low levels of activity for both groups (**Figure 4C**). At days 35 and 42, however, IEE iso-CTL neurons exhibited a 1.49 Hz increase in weighted mean firing rate (WMFR) (**Figure 4D**, p = 0.01) and an 0.95 Hz increase in WMFR (**Figure 4E**, p = 0.04), respectively. By contrast, ↑GIRK2 neurons appeared unaffected by two weeks of IEE at day 35 (**Figure 4C**), but were significantly inhibited after three weeks of IEE at day 42, with a statistically significant interaction effect between increased GIRK2 expression and ethanol exposure (−1.37 Hz WMFR; p = 0.02) (**Figure 4E**).

### GIRK2-upregulated neurons show inhibited spontaneous calcium spiking activity after 21-day IEE

To investigate the combined effects of 21-day IEE and ↑GIRK2 on glutamatergic neuronal activity under ethanol-free conditions, we conducted Ca^2+^ imaging, using Ca^2+^ transients as a proxy for neuronal activity. We expanded our donor pool to five cell lines (CRISPRa: 553/dCas9-VPR and 2607/dCas9-VPR; Lenti: 12455, 9429, and BJ) and carried out the same IEE protocol with *NGN2-*induced glutamatergic neurons, as described for the MEA experiments (**Figure 4A**). Ca^2+^ transients were detected with Fluo4-AM, and were baseline-corrected, filtered for non-neuronal events, and de-convolved to quantify peak frequency and height (**Figure 5A-C**) (N = 50-200 regions of interest (ROI)|5 donors|3 replicates; see extended data Table 5-1 for all ROI N values for each replicate experiment). Neurons without spontaneous activity were excluded from spontaneous activity analysis.

**Figure 5.**
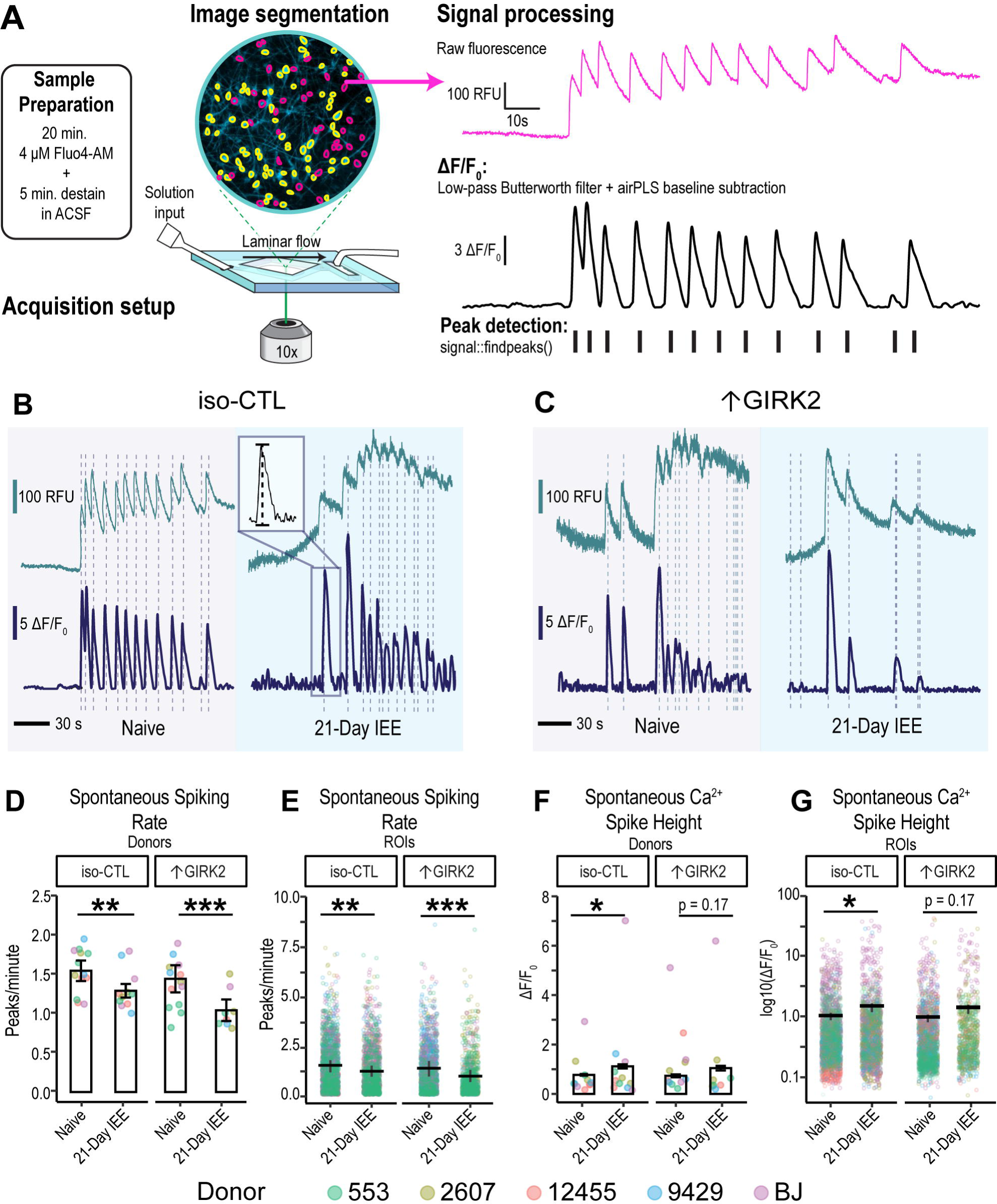
Combined effect of 21-Day IEE and ↑GIRK2 expression inhibits spontaneous activity of excitatory neurons. **A.** Schematic representation of calcium imaging experimental and analysis procedure, including sample preparation, acquisition setup, and signal processing. **B-D:** Representative fluorescence traces showing Ca^2+^ spikes in alcohol-naïve (grey background) and 21-day IEE-treated (blue background) iso-CTL **(B)** and ↑GIRK2 **(C)** excitatory neurons on D42. Inset in iso-CTL: zoom of Ca^2+^ spike illustrating peak height measurement. Vertical dashed lines indicate auto detection of Ca^2+^ peaks. Top trace is raw fluorescence. Bottom trace is baseline-corrected ΔF/F_0_. Scale bars = 100 RFU, 5 ΔF/F_0_, and 30s. See Extended Data Figure 4-1 for schematic of Ca^2+^ fluorescence acquisition and processing pipeline. **D, E**: Ca^2+^ spike rate (peaks/minute) plotted as an average for each donor replicate (n = 5 donors, 3 replicates) **(D)** and as individual neuronal ROIs (see Extended Data Table 5-1 for complete list of spontaneously active neuronal ROI Ns for each replicate experiment) **(E)**. **F-G:** Ca^2+^ transient height (ΔF/F_0_) plotted as an average for each donor replicate **(F)** and individual ROIs **(G)**. Color indicates donor (coral = 12455; dark yellow = 3440; green = 553; blue = 9429; and purple = BJ). GLMM includes replicate and donor random effects. * p < 0.05, ** p < 0.01, *** p < 0.001.

Similar to MEA findings, IEE for 21 days significantly reduced the frequency of calcium spiking in spontaneously active neurons in ↑GIRK2 neurons in ethanol-free solution (decrease of –0.51 peaks/minute, p < 0.00001 IEE effect in ↑GIRK2 neurons; p = 0.0006 interaction ↑GIRK2:IEE) (**Figure 5D-E**). One interpretation of this finding is that the inhibitory interaction of IEE with ↑GIRK2 is not caused by a direct interaction of ethanol on GIRK2 channels. In contrast to MEA results, iso-CTL neurons exhibited decreased spontaneous Ca^2+^ activity as a result of IEE under ethanol-free conditions, albeit to a lesser extent than ↑GIRK2 neurons (–0.19 peaks/minute, p = 0.003) (**Figure 5D-E**). IEE iso-CTL neurons also displayed a significant increase in calcium transient peak heights (+0.33 ΔF/F_0_, p =0.002) (**Figure 5F-G**), possibly indicating stronger depolarization. ↑GIRK2 neurons, on the other hand, showed no significant change in Ca^2+^ levels (+0.16 ΔF/F_0_, p = 0.10), suggesting that ↑GIRK2 expression ablates ethanol-induced increases in intracellular Ca^2+^.

### GIRK2 expression interacts with ethanol to differentially regulate glutamate response and intrinsic excitability

We hypothesized that, in addition to inhibiting spontaneous activity, 21-day ethanol exposure would alter neuronal excitability in ↑GIRK2 neurons. We interrogated neuronal excitability by rapid bath application of glutamate (10 μM for 30 s), which would be expected to depolarize neurons and elicit Ca^2+^ spikes. We measured the proportion of glutamate-responsive neurons, the number of Ca^2+^ peaks, the latency to the first spike, and Ca^2+^ peak height (**Figure 6C**). In iso-CTL neurons, 21-day IEE increased the proportion of glutamate-responsive neurons (+13.3%, p < 0 .001; N = 1203-2609 ROIs/group; see extended data **Table 6-1** for full list of ROI N values for each donor & replicate) (**Figure 6D**). In the glutamate-active subset of neurons, IEE also increased the number of glutamate-evoked spikes (+ 0.2 spikes, p = 0.02) (**Figure 6A, E**). These effects of IEE were absent in ↑GIRK2 neurons (**Figure 6B, D-E**). Furthermore, IEE decreased latency to first spike with glutamate (**Figure 6C**) in iso-CTL neurons, suggesting an increase in response to glutamate and/or excitability (–3.4 s, p < 0.00001) (**Figure 6F**). By contrast, 21-day IEE ↑GIRK2 neurons increased their latency to first spike with glutamate (+2.1 s, p = 0.0312 exposure effect; p < 0.00001 interaction effect) (**Figure 5F**).

**Figure 6.**
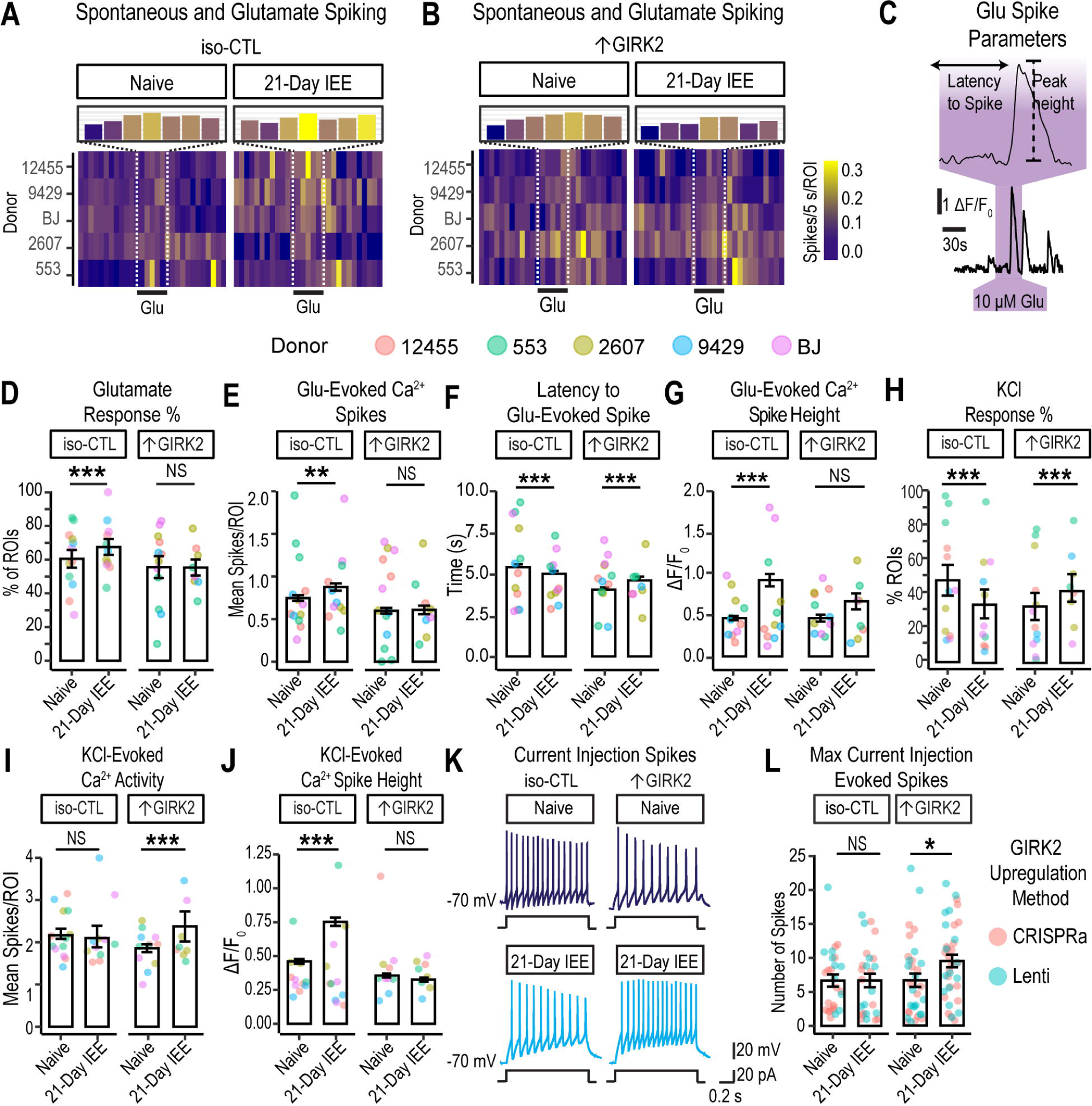
Increased GIRK2 expression and intermittent ethanol exposure impact glutamate response and membrane excitability in glutamatergic neurons. **A, B.** Calcium spiking activity heatmap for iso-CTL **(A)** and ↑GIRK2 **(B)** neurons, either naive and with 21-day IEE. 5-second bins, normalized to number of cells (spikes/5 s/ROI). Inset magnification shows number calcium spikes/5 s/ROI during the 30 s pulse of 10 μM glutamate. Color gradient indicates activity intensity (violet-blue = low; yellow = high). **C.** Example deconvolved Ca^2+^ fluorescence trace, and latency to spike with a 30 s glutamate pulse. **D**. % of excitatory neurons responsive to 30 s glutamate pulse (10 µM). (See Extended Data Table 6-1 for complete list of all active neuronal ROI Ns for each replicate experiment). **E.** Number of calcium spikes during each 30s glutamate pulse (See Extended Data Table 6-2 for complete list of all glutamate-responsive ROI Ns for each replicate experiment). **F**. Latency to spike (s) in response to 30 s glutamate, pulse (10 µM). **G.** Calcium peak height (ΔF/F_0_) evoked by glutamate for iso-CTL and ↑GIRK2 neurons, either ethanol-naive or with 21-day IEE. **H.** % of ROIs responsive to 15 mM KCl. **I.** Number of spikes evoked by 1 min 15 mM KCl pulse. **J.** Ca^2+^ spike height during 1 min 15 mM KCl (ΔF/F_0_). KCl. (N = 1203-2608 ROIs/group). Point color indicates donor (coral = 12455; dark yellow = 2607; green = 553; blue = 9429; and purple = BJ). (See Extended Data Table 6-3 for complete list of glutamate-responsive and KCl-active ROI Ns for each replicate experiment). **K.** Representative action potentials evoked by 1s current injection step (20 pA) in naïve and 21-day IEE iso-CTL and ↑GIRK2 neurons. **L.** Maximum number of evoked spikes measured in current-clamp with 20 pA current injection steps from holding potential of ∼ −70 mV (N = iso-CTL: 26 Naïve, 25 21-day IEE; ↑GIRK2: 33 Naïve, 37 21-Day IEE). Point color indicates upregulation method (coral = CRISPRa; teal = Lenti). GLMM includes batch and donor effects. * p < 0.05, ** p < 0.01, *** p < 0.001.

To further validate of latency calculation, we examined the latency to the first spike with a random 30s epoch in the same neuron. The average latency to the first spike was 14.7 ± 0.3, 15.1 ± 0.3, 15.2 ± 0.3, and 16.0 ± 0.4 for naïve iso-CTL, 21-day IEE iso-CTL, naïve-↑GIRK2, and 21-day IEE-↑GIRK2 groups, respectively; p values equal to 0.92 and 0.39 for the impact of IEE on iso-CTL and ↑GIRK2 neurons, respectively. By contrast, the latency to the first spike seen with glutamate application was 4.7 ± 0.2, 4.6 ± 0.2, 4.1 ± 0.2, and 4.4 ± 0.4 for each treatment group, as listed above, confirming the latency to the first spike reflects a response to glutamate.

Similar to the measurements of spontaneous activity (**Figure 5F-G**), calcium peak heights were significantly larger in IEE iso-CTL neurons (+0.36 ΔF/F_0_, p = 3×10^-6^). This increase was not observed in ↑GIRK2 neurons (**Figure 6G**). Taken together, these data indicate that 21-day IEE heightens glutamate receptor activity and/or excitability in iso-CTL neurons, whereas ↑GIRK2 neurons exhibit diminished glutamate responses.

To determine whether differences in glutamate-responsive neurons were due to changes in glutamate sensitivity or in intrinsic excitability, we bath-applied 15 mM KCl in ACSF to directly depolarize the neurons (15 mM KCl is predicted to produce a ∼32 mV positive shift in the resting membrane potential) and measured Ca^2+^ spikes in the same population of glutamate-responsive neurons. As before, Ca^2+^ flux was elevated in 21-day IEE iso-CTL neurons (+ 0.23 ΔF/F_0_, p < 1×10^-6^), while ↑GIRK2 neurons were unaffected (**Figure 5J**). However, 21-day IEE in ↑GIRK2 neurons increased proportion of KCl-responsive ROIs (+8.3%, p = 0.009 IEE effect; p < 1×10^-6^ interaction effect) (**Figure 5H**) and number KCl-evoked Ca^2+^ spikes (+ 0.54 spikes, p < 1×10^-6^) (**Figure 5I**). Iso-CTL neurons, conversely, had a lower proportion of KCl-responsive ROIs (−16.5%, p < 1X10^6^) (**Figure 5H**) and no difference in number of evoked spikes (p = 0.4) (**Figure 5I**). These data indicate that, unlike glutamate-evoked activity, depolarization-evoked activity is suppressed in iso-CTL neurons and heightened in ↑GIRK2 neurons following 21-day IEE.

To further probe potential differences in intrinsic excitability, we examined the maximum number of spikes induced with depolarizing current injection steps (20 pA) using whole-cell patch-clamp electrophysiology (**Figure 6K**). Similar to KCl-induced firing (**Figure 6I**), there was no significant difference in the number of spikes for iso-CTL neurons with 21-day IEE. By contrast, we recorded a higher number of spikes in ↑GIRK2 neurons with 21-day IEE (+2.8 spikes, p = 0.002) (**Figure 5K-L**), similar to that with KCl-induced firing. Taken together, these results suggest inhibitory effects of ↑GIRK2 with 21-day IEE appear to be specific to glutamate-mediated activity.

To investigate potential molecular mechanisms underlying combined effects of increased GIRK2 expression and intermittent ethanol exposure on glutamatergic function, we performed bulk RNAseq differential gene expression analysis in the same cohort of neurons subject to 21-day IEE. Acting in concert, ↑GIRK2 and IEE impacted expression of multiple genes essential for synaptic function (**Figure 7, Extended Data Table 7-1**). In contrast to untreated isogenic controls, neurons treated with IEE +↑GIRK2 exhibited down-regulation of glutamatergic receptor (*GRIA1, GRIN1* & *GRIN2B*) and potassium channel (*KCND2* & *KCNJ3*) genes (**Figure 7A**). This transcriptomic landscape is consistent with the decreased response to glutamate and increased membrane excitability in IEE + ↑GIRK2 neurons (**Figure 6**). Furthermore, the combination of IEE and ↑GIRK2 led to downregulation of Ca^2+^ signaling genes, including the voltage-gated Ca^2+^ channel *CACNAD1* and intracellular Ca^2+^-binding protein *CALB2* (**Figure 7A**). Reduced expression of Ca^2+^- sensitive genes may reflect an active process in ↑GIRK2 neurons working in opposition to IEE-induced increases in intracellular calcium present in isogenic controls both during spontaneous (**Figure 5**) and evoked (**Figure 6**) neuronal activity.

**Figure 7.**
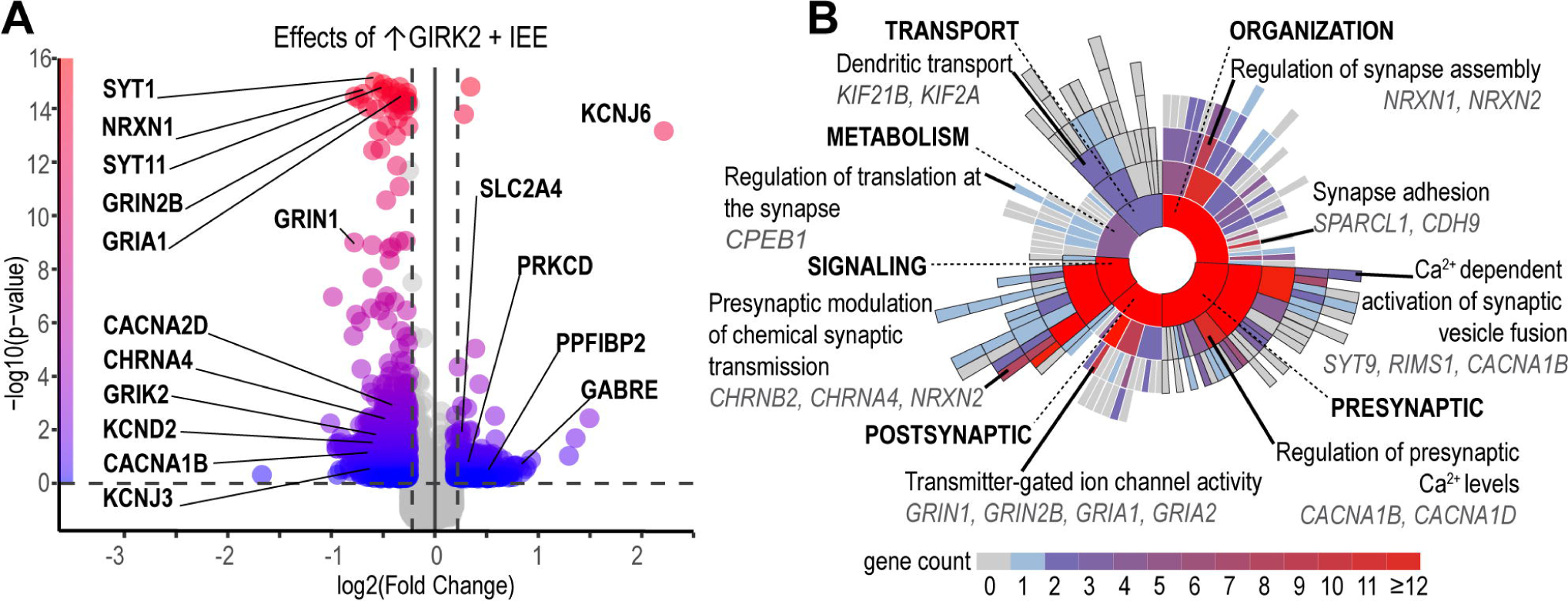
Intermittent ethanol 21-day exposure combined with↑GIRK2 expression uniquely impacts synaptic function gene pathways. **A.** Volcano plot of differentially regulated genes (1.25 log2 fold change and 0.05 p-value cutoffs) as a result of the combined effect of IEE and ↑GIRK2 on iso-CTL neurons. Color gradient indicates -log10(p-value), with blue = 0 and red = 16. Select neuronal excitability genes highlighted – see Extended Data Table 7-1 for full list of differentially regulated synaptic function transcripts resulting from combined**↑**GIRK2 and 21-day IEE. **B.** Sunburst plot of SynGO annotated gene ontologies, including synaptic transport, metabolism, organization, signaling, presynaptic and post-synaptic biological processes. Color indicates number of genes enriched in each pathway, including child terms (grey = 0, blue = 1, red ≥ 12). Top enriched pathways and differentially expressed genes are highlighted for each biological process. See Extended Data Table 7-2 for complete list of annotated SynGO pathways and their enrichment values. N = 2 donors CRISPRa, 4 donors Lenti | 2 batches | 3 technical replicates/donor/batch.

To gain insight into how these shifts in RNA expression may contribute to specific biological processes involved in neuronal signaling, we analyzed synaptic gene ontology using the SynGO online portal (Koopmans et al., 2019) (**Figure 7B, Extended Data Table 7-2**). While each of the main biological processes was significantly impacted by combined ↑GIRK2 and IEE (i.e., pre- and post-synaptic function, signaling, transport, organization, and metabolism), pre-synaptic pathways were the most affected (**Fig. 7**). Specifically, regulation of presynaptic Ca^2+^ levels and Ca^2+^-dependent vesicle fusion (both involving *CACNA1B*) appeared especially sensitive to combined ↑GIRK2 and IEE (**Fig. 7**). These alterations in presynaptic regulatory mechanisms occurred in conjunction with changes in synapse assembly, synaptic translation, and dendritic transport. Taken together, these data support an active transcriptomic process underlying the observed changes in neuronal activity (e.g., altered glutamate response, intrinsic excitability, and intracellular calcium handling) resulting from the dual effects of ↑GIRK2 and IEE.

### Impact of ethanol exposure, GIRK2 expression, and glutamate on mitochondrial function in excitatory neurons

SynGO analysis (**Figure 7B, Extended Data Table 7-2**) indicated adaptations in metabolic processes involved in translation at the synapse – an integral aspect of activity-dependent modulation of neuronal circuitry. Furthermore, transcriptomic analysis in day 28 neurons following 7-day IEE (**Figure 3D**) suggested that metabolic and mitochondrial alterations is an early response to ethanol, preceding significant changes to glutamatergic signaling. Together, these suggested a role for mitochondria with chronic alcohol use (Vandecasteele et al., 2001; Cassano et al., 2016; Giorgi et al., 2018; Verma et al., 2022).

We therefore investigated the effect of IEE and ↑GIRK2 on the developmental time-course of mitochondrial function in glutamatergic neurons. We hypothesized that ↑GIRK2 would prevent changes in cellular respiration caused by 7-day and 21-day IEE, similar to the mitigating effect on increased calcium activity. To test this prediction, we utilized the mitochondrial stress test (Agilent Technologies Seahorse assay) with monocultures of glutamatergic neurons (monocultures eliminate the potential confound of glial cells). Respiration was measured in ethanol-free conditions at day 28 post-induction with 7-day IEE and at day 42 with 21-day IEE (N = 5 donors|4 wells|3 batches/timepoint) and normalized for cell density. Seven-day IEE resulted in significant impairment of basal (–17.5 pmol/min/cell oxygen consumption rate (OCR), p = 0.03) (**Figure 8A** and **8C**) and maximal (–72.6 pmol/min/cell OCR, p = 0.04) (**Figure 8A** and **8D**) cellular respiration in iso-CTL neurons. By contrast, ↑GIRK2 neurons appeared less sensitive to IEE (–10.5 pmol/min/cell OCR, p = 0.61). This decrease in respiration in IEE iso-CTL neurons may be the stimulus for upregulation of mitochondrial and metabolic genes as measured in RNAseq (**Figure 3D**), serving as a potential compensatory mechanism.

**Figure 8.**
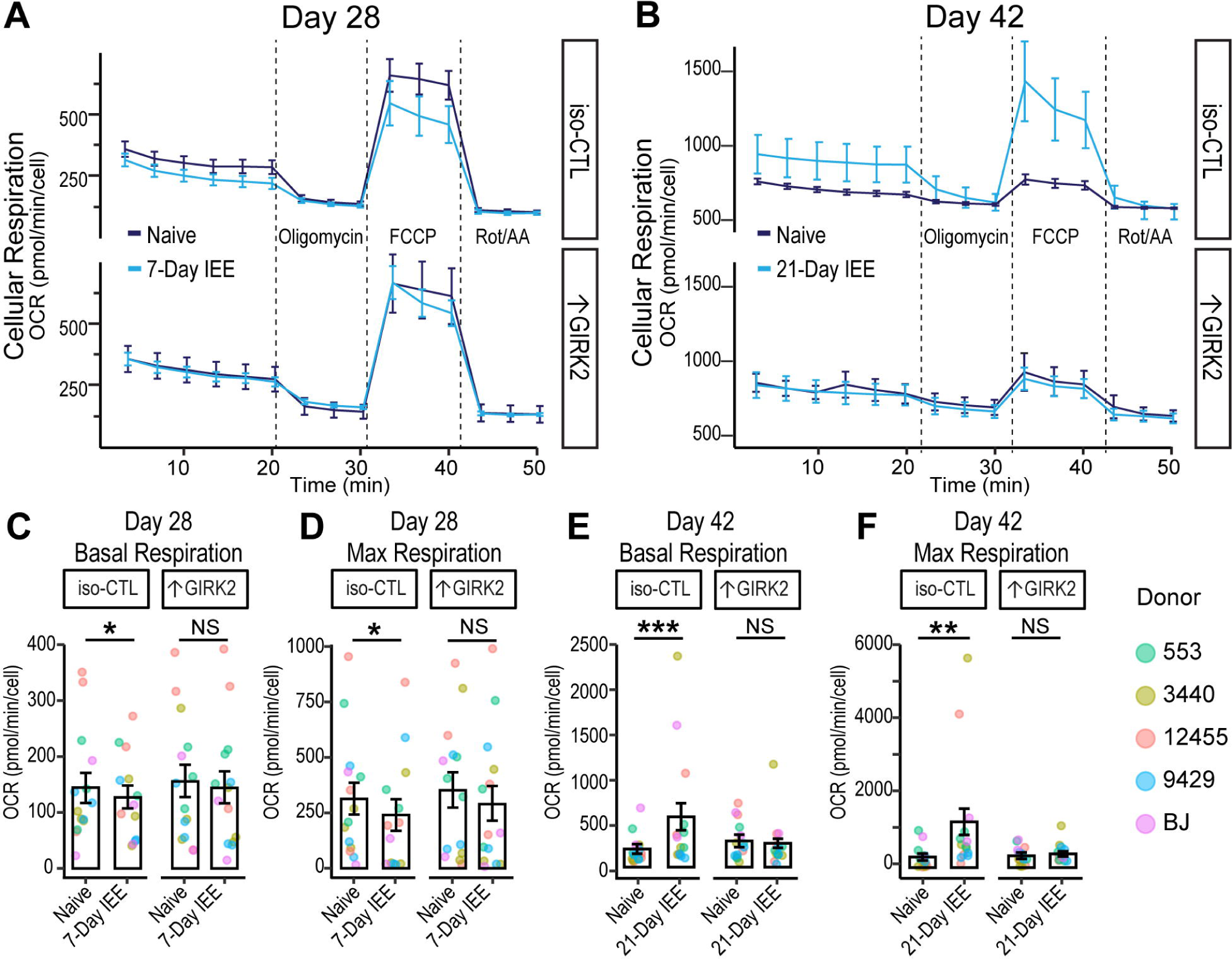
↑GIRK2 expression counteracts effects of IEE on mitochondrial function. **A-B:** Time course of cell-density-normalized oxygen consumption rate (OCR; pmol/minute/cell) during the mitochondrial stress test in CTL and ↑GIRK2 neurons at Day 28 (7-day IEE) **(A)** and Day 42 (21-day IEE) **(B)** post-induction. Line color indicates ethanol exposure: dark blue = naive, light blue = IEE. **C-F:** Day 28 post-induction basal **(C)** and maximal OCR (pmol/min/cell) **(D)** and D42 basal **(E)** and maximal **(F)** OCR in WT and ↑GIRK2 neurons naive to ethanol or with IEE. * p < 0.05, **p < 0.01, *** p < 0.001.

At day 42, the overall oxygen consumption rate was higher than at day 28 (**Figure 8A-F**), reflecting a parallel between increased neuronal activity (**Figure 4C-E**) and increased oxygen consumption. Twenty-one-day IEE significantly increased oxygen consumption during basal (+352 pmol/min/cell OCR, p = 0.0006) (**Figure 8E**) and maximal (+928 pmol/min/cell OCR, p = 0.00003) (**Figure 8F**) respiration in iso-CTL neurons. Notably, ↑GIRK2 neurons did not show the increase in basal or maximal respiration with 21-day IEE (**Figure 8B**). Statistical analysis indicates an interaction between ↑GIRK2 and ethanol exposure for both basal (p = 0.02) (**Figure 8E**) and maximal (p = 0.009) (**Figure 8F**) respiration. These results suggest a degree of resilience and potential pro-allostatic influence from ↑GIRK2.

To test whether these changes were reflective of activity-dependent energy demands, we stimulated neuronal activity with an acute injection of 10 μM glutamate and measured ATP-linked respiration (**Figure 9A-B**). At day 28, we observed no differences in ATP-linked respiration at basal levels of activity, regardless of ethanol exposure or GIRK2 expression levels (**Figure 9C**). The glutamate (10 μM) challenge, on the other hand, reduced ATP-linked respiration in IEE iso-CTL (–66.3 pmol/min/cell OCR, p = 0.017) but not in ↑GIRK2 neurons (p = 0.49) (**Figure 9C**). This suggests an initial impairment in the ability of iso-CTL neurons to meet activity-dependent energy demands, which are likely lessened by ↑GIRK2 expression.

**Figure 9.**
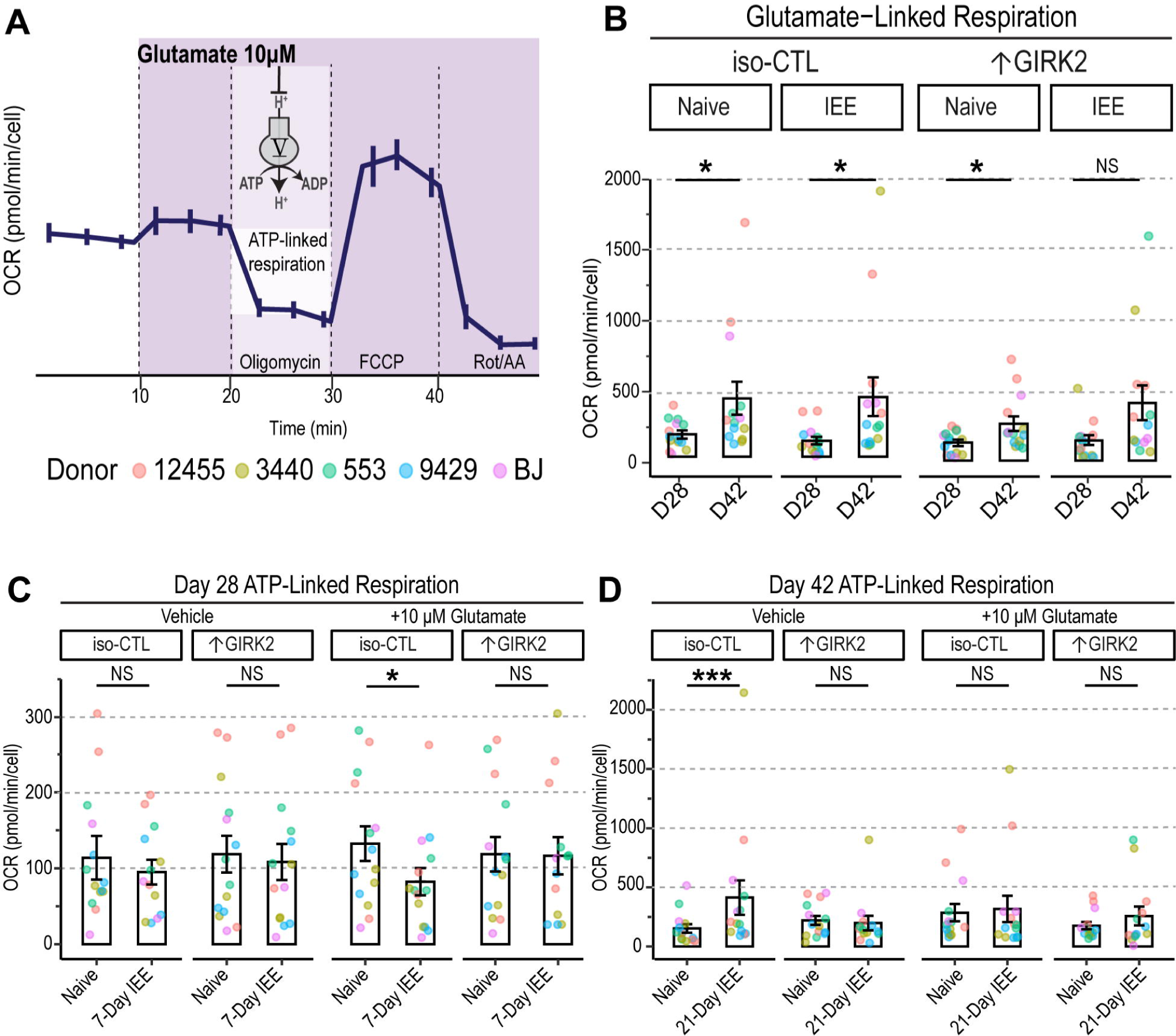
↑GIRK2 expression counteracts IEE-induced deficit in ATP production in the presence of glutamate. **A.** Schematic representation of acute 10 µM glutamate injection during the mitochondrial stress test and mechanism of ATP-linked respiration measurement. **B.** Glutamate linked respiration in iso-CTL and ↑GIRK2 *NGN2* neurons at 28- and 42-days post-induction (D28 and D42, respectively). Each point represents an average of 4 wells from 3 replicate experiments (N = 5 donors). **C-D:** ATP linked respiration in D28 **(C)** and D42 **(D**) neuron naive to ethanol or with IEE during basal (vehicle) and stimulated (10 μM glutamate) conditions (pmol/min/cell). Each point represents and average of 4 wells from 3 replicate experiments (N = 5 donors). Point color indicates donor (coral = 12455; dark yellow = 3440; green = 553; blue = 9429; purple = BJ). GLMM includes donor and batch effects. * p < 0.05, **p < 0.01, *** p < 0.001.

At day 42, iso-CTL neurons exhibited a surge in ATP-linked respiration (+260 pmol/min/cell OCR, p = 0.03) (**Figure 9B, D**) under basal conditions, consistent with the observed increases in basal (**Figure 8E**) and maximal (**Figure 8F**) oxygen consumption rate. Interestingly, the glutamate (10 μM) challenge eliminated these effects, boosting respiration in ethanol-naïve neurons and dampening respiration in ethanol-exposed iso-CTL neurons. Since ethanol exposure increases glutamate sensitivity and firing in iso-CTL neurons, these findings support the conclusion of increased energy demands and the involvement of compensatory mechanisms. In line with the reduced glutamate response evident in Ca^2+^ imaging (**Figure 6B, D-G**), ATP-linked respiration remained unchanged in ↑GIRK2 neurons following 21-day IEE (p = 0.57) (**Figure 9B, D**).

Overall, these transcriptomic and functional data support our observations of increased GIRK2 moderating the IEE-induced mitochondrial respiratory alterations. Mitochondrial stress tests data represent a progression from an initial impairment to a subsequent surge of mitochondrial function as a result of prolonged ethanol exposure. Our findings also suggest that ethanol exposure confers a metabolic challenge, which is exacerbated by the presence of elevated glutamate. Increased GIRK2 expression appears to prevent this trajectory, as evidenced by unaltered cellular respiration regardless of ethanol exposure or glutamate challenge. Further investigation of mitochondrial localization, integrity, and neuronal metabolome is needed to parse the precise role of increased GIRK2 on metabolic adaptations to ethanol exposure in excitatory neurons.

## DISCUSSION

In the current study, we employed an isogenic approach to directly examine the functional effects of GIRK2 channels in human excitatory glutamatergic neurons, and to investigate their role in adaptations to physiological concentrations of ethanol over 21 days. We hypothesized that increases in GIRK2 expression would impact neuronal response to ethanol exposure, either through direct interaction with the alcohol-binding pocket in GIRK2, and/or through secondary mechanisms and regulation of neuronal excitability. We found that increased GIRK2 expression (↑GIRK2) mitigates changes in neuronal activity and mitochondrial respiration induced by 21-day intermittent 17 mM ethanol exposure.

Human neurons with ↑GIRK2 exhibited more hyperpolarized resting potentials, and larger GIRK-like currents (SCH-inhibited), as expected for expression of an inwardly-rectifying potassium channel (Kuzhikandathil and Oxford, 2002; Lüscher and Slesinger, 2010; Zhao et al., 2020). A major finding was that several of the changes in spontaneous neuronal activity induced by IEE (including increased MEA firing and increased amplitude of Ca^2+^ transients) were absent in ↑GIRK2 neurons, suggesting a mitigating effect of increased GIRK2. We explored whether these changes occurred through differences in glutamate response or intrinsic excitability, uncovering changes in both excitability and suppression of IEE-induced increase in glutamate sensitivity with ↑GIRK2 expression. The increase in neuronal activity with glutamate (i.e., increased % of responsive neurons, decreased latency to spike, and increased number of spikes/glutamate pulse) following IEE was blunted by ↑GIRK2.

Transcriptomic data suggested that IEE-induced increase in glutamate responsiveness for iso-CTL neurons was not attributable to changes in glutamate receptor expression, implicating processes downstream of RNA, such as phosphorylation and ubiquitination regulation of receptor trafficking, surface expression, and turnover (Desch et al., 2021; Warner et al., 2022). Combination of 21-day IEE with ↑GIRK2 neurons reduced expression of NMDAR subunits 1 and 2B and AMPAR subunit 1, possibly reflecting transcriptomic compensation for altered glutamate sensitivity. Furthermore, synaptic gene ontology analysis indicated a role for GIRK2 in presynaptic mechanisms contributing to glutamatergic adaptations to IEE. Specifically, differential regulation of voltage-gated Ca^2+^ channel subunits (including *CACNA1B, CACNA2D*) and synaptotagmin family members (including *SYT1, 4, 5, 7* and *11*) points to calcium-mediated vesicle release as a major consequence of combined ↑GIRK2 and IEE. Additional experiments, such as measuring miniature EPSCs and synaptosome proteomics, are necessary to assess the possible presynaptic changes mediated by IEE and ↑GIRK2 interactions.

Unexpectedly, ↑GIRK2 neurons appeared to be more excitable with depolarization following IEE. This change could result from the decreased potassium channel expression seen in RNAseq data. Alternatively, a reduced voltage-dependent inactivation of Na^+^ as a consequence of hyperpolarization in ↑GIRK2 neurons with IEE could also increase neuronal excitability (Bähring et al., 2012; Fourcaud-Trocmé et al., 2022; Ransdell et al., 2022). Together, these findings favor the conclusion that the synergistic effect of IEE and ↑GIRK2 expression targets glutamate-receptor-dependent signaling. Further investigation is necessary to ascertain the relationship between transcriptomic responses to IEE and corresponding states of protein activity.

Increased GIRK2 expression also prevented increases in the amplitude of calcium peaks induced by IEE in iso-CTL neurons. Intracellular calcium dynamics are multiplex, and the amplitude of calcium peaks could be affected by neuronal firing activity and/or changes in voltage-gated calcium channels. Since elevated intracellular calcium is a hallmark of excitotoxicity (Verma et al., 2022), GIRK2-mediated inhibition in ethanol-exposed glutamatergic neurons may serve a protective role. Consistent with this interpretation, calcium influx through NMDAR channels has been shown to increase surface expression of GIRK channels (Chung et al., 2009), supporting the conclusion that GIRK channel expression is linked to fluctuations in glutamatergic signaling. GIRK channels are also important for synaptic depotentiation (Chung et al., 2009).

In support of our finding that the upregulation of GIRK2 plays an important role in modulating excitability and response to ethanol in human glutamatergic neurons, a recent study found that neurons derived from AUD-diagnosed participants with specific *KCNJ6* SNPs had lower levels of GIRK2 expression, greater neurite area, and heightened excitability as compared to neurons derived from unaffected individuals (Popova et al., 2023). Overexpression of GIRK2 in neurons from AUD-diagnosed individuals mimicked the effects of the 7-day IEE protocol, eliminating differences in neuronal morphology induced excitability (Popova et al., 2023). Thus, in the context of AUD, increased GIRK2 expression and subsequent neuronal inhibition appears to exert a normalizing influence, possibly through regulatory mechanisms of neurotransmitter release and intracellular calcium levels.

Given the impact of increased GIRK2 on intracellular calcium in neurons exposed to ethanol, ↑GIRK2 may also help maintain normal mitochondrial function. Recent work by Sun *et al*. showed that glutamatergic neurons metabolize ethanol, using it as a preferred energy source over glucose in chronic ethanol exposure conditions (Sun et al., 2023). Mitochondria are vulnerable to ethanol-induced metabolic changes (Tapia-Rojas et al., 2017; León et al., 2022; Lim et al., 2023) and their regulation of calcium homeostasis plays a key role in excitotoxicity (Vandecasteele et al., 2001; Giorgi et al., 2018; Verma et al., 2022). The upregulation of mitochondrial and metabolic genes after 7-day IEE suggests an early metabolic response to ethanol. Furthermore, several groups reported that ethanol exposure of iPSC-derived neurons affected mitochondrial health and neuronal development (Gunnewiek et al., 2019; Motori et al., 2020), including upregulation of genes involved in cholesterol homeostasis (*INSIG1* and *LDLR*) (Jensen et al., 2019). Although the ethanol concentration used varied (17 mM in our study vs. 50 mM in Jensen et al., 2019), the results are similar. The top two upregulated genes in our 7-day ethanol exposure paradigm (*ANGPTL4* and *PDK4*) (Figure 3B) are also key players in lipid metabolism; *PDK4* in particular, has been associated with a loss of metabolic flexibility (Zhang et al., 2014; Jeon et al., 2020) and disrupted mitochondrial homeostasis (Ma et al., 2019). Notably, GIRK channel activity is potentiated by cholesterol (Glaaser and Slesinger, 2017; Mathiharan et al., 2020), suggesting that changes in membrane cholesterol could impact GIRK function.

In parallel with maintaining intracellular calcium dynamics, ↑GIRK2 neurons appeared to preserve normal mitochondrial function, regardless of ethanol exposure (7- and 21-day IEE) or activity-dependent energy demands. By contrast, stimulation of neuronal activity in iso-CTL neurons with glutamate revealed an impairment in ATP-linked respiration after 7-day IEE, indicating a difficulty in meeting activity-dependent energy demands. Furthermore, after 21-day IEE, glutamate eliminated the increase of ATP-linked respiration in iso-CTL neurons. The reduction of ATP-linked respiration with the application of glutamate may be attributable to increased calcium influx and the disruption of mitochondrial membrane potential (Giorgi et al., 2018; Verma et al., 2022). SynGO analysis of combined ↑GIRK2 and IEE points to regulation of synaptic translation as a potential mechanism underlying the apparent preservation of normal mitochondrial function.

Previous studies investigating the role of GIRK2 expression in neuronal health showed somewhat conflicting outcomes. One study demonstrated that GIRK2 expression in TH-positive substantia nigra dopaminergic neurons is a vulnerability factor to mitochondrial stress and apoptosis (Chung et al., 2005). Another found that GIRK2 is upregulated in response to neurotoxins such as Aβ_42_ in hippocampal dissociated cultures, potentially triggering p75^NTR^ mediated cell-death (May et al., 2017). Notably, Aβ_42_ has an excitotoxic effect similar to the consequences of long-term ethanol exposure (Esposito et al., 2013), and therefore GIRK2 upregulation could be a protective measure. Further studies, including mitochondrial imaging and viability assays, are required to shed light on mechanisms underlying the relationship between GIRK2 expression, presynaptic calcium-dependent regulation of neurotransmitter release, synaptic translation, and mitochondrial function.

### Future Perspectives

In the current study, we focused on glutamatergic neurons due to their primary role in addiction (Burnett et al., 2016). Given the broad expression of GIRK channels as well as the regional and cell-type variability of their impact on neuronal activity, however, our observation could be unique to maturing glutamatergic neurons in the absence of inhibitory control. Future work should investigate mono- and co-cultures with GABAergic and dopaminergic neurons, given the role of GIRK channels in facilitating GABA_B_ and D_2_ receptor function (Lüscher and Slesinger, 2010). Furthermore, 3D models such as assembloids and organoids can shed light on the role of GIRK2 and ethanol exposure on circuit development and neurodegenerative processes (Birey et al., 2017; Arzua et al., 2020; Szebényi et al., 2021). Differences in open state probability between GIRK2 homotetramer and GIRK1/2 heterotetramer channels (Rubinstein et al., 2009) also could play a role in neuronal excitability. Together, these future directions pave a path toward a cell- and circuit-level understanding of GIRK2-containing channels in human neurons and their influence on developmental and apoptotic processes affected by ethanol exposure.

### Conclusions

In summary, we demonstrate that GIRK2 expression impacts the long-term effects of ethanol on neuronal activity and bioenergetics of glutamate neurons, suggesting a possible role for GIRK2 in mitigating dysfunctions caused by chronic alcohol use. Further investigations are necessary to determine the mechanisms underlying the intersection of GIRK channels, metabolism, and neuronal excitability and how they may vary in different neuronal cell types. Several key questions remain, including whether basal GIRK channel activity is essential for preserving mitochondrial function and whether increasing GIRK2 expression after the start of ethanol exposure would have a similar ameliorating effect. The relationship between GIRK2 activity and cellular respiration represents a potential new link with alcohol, opening a novel line of inquiry into the channel’s role in neuronal health and resilience in the context of AUD. Initial levels of GIRK2 play a role in neuronal adaptations to ethanol and could therefore serve as predictors of pharmacotherapy response, allowing for the development of personalized treatments for AUD and prenatal alcohol exposure.

## ACKNOWLEDGMENTS

We gratefully acknowledge the support provided by the Ruth L. Kirschstein National Research Service Award (NRSA) Individual Predoctoral Fellowship (1 F31AA027949-01), and Collaborative Studies on the Genetics of Alcoholism (National Institute on Alcohol Abuse and Alcoholism; NIAAA U10AA008401). Special thanks to Dr. Kristen Brennand and Dr. Julia TCW for providing NPC cell lines used in this study. Additional thanks to Dr. Edoardo Marcora for consultation on generalized linear mixed model statistical analysis.

The Collaborative Study on the Genetics of Alcoholism (COGA), Principal Investigators B. Porjesz, V. Hesselbrock, T. Foroud; Scientific Director, A. Agrawal; Translational Director, D. Dick, includes eleven different centers: University of Connecticut (V. Hesselbrock); Indiana University (H.J. Edenberg, T. Foroud, Y. Liu, M. Plawecki); University of Iowa Carver College of Medicine (S. Kuperman, J. Kramer); SUNY Downstate Health Sciences University (B. Porjesz, J. Meyers, C. Kamarajan, A. Pandey); Washington University in St. Louis (L. Bierut, J. Rice, K. Bucholz, A. Agrawal); University of California at San Diego (M. Schuckit); Rutgers University (J. Tischfield, R. Hart, J. Salvatore); The Children’s Hospital of Philadelphia, University of Pennsylvania (L. Almasy); Virginia Commonwealth University (D. Dick); Icahn School of Medicine at Mount Sinai (A. Goate, P. Slesinger); and Howard University (D. Scott). Other COGA collaborators include: L. Bauer (University of Connecticut); J. Nurnberger Jr., L. Wetherill, X., Xuei, D. Lai, S. O’Connor, (Indiana University); G. Chan (University of Iowa; University of Connecticut); D.B. Chorlian, J. Zhang, P. Barr, S. Kinreich, G. Pandey (SUNY Downstate); N. Mullins (Icahn School of Medicine at Mount Sinai); A. Anokhin, S. Hartz, E. Johnson, V. McCutcheon, S. Saccone (Washington University); J. Moore, Z. Pang, S. Kuo (Rutgers University); A. Merikangas (The Children’s Hospital of Philadelphia and University of Pennsylvania); F. Aliev (Virginia Commonwealth University); H. Chin and A. Parsian are the NIAAA Staff Collaborators. We continue to be inspired by our memories of Henri Begleiter and Theodore Reich, founding PI and Co-PI of COGA, and also owe a debt of gratitude to other past organizers of COGA, including Ting-Kai Li, P. Michael Conneally, Raymond Crowe, and Wendy Reich, for their critical contributions. This national collaborative study is supported by NIH Grant U10AA008401 from the National Institute on Alcohol Abuse and Alcoholism (NIAAA) and the National Institute on Drug Abuse (NIDA).

## AUTHOR CONTRIBUTIONS

IP, AG, PAS conceived the project. IP and MBF performed experiments. IP and YL performed statistical analyses and discussed with PAS and AG. AG, PAS, RPH, ZPP, IGR, DP provided critical feedback throughout the project. IP, PAS prepared the paper with critical evaluation from RPH, ZPP, JST, BP, HJE, XX, DPC, CK. CK, RPH, HJE, and BP edited the manuscript.

