## Extended Data Table 5-1 for "Upregulated GIRK2 counteracts ethanol-induced changes in excitability & respiration in human neurons"

**Extended Data Table 5-1: Spontaneously active iso-CTL and ↑GIRK2 neurons naïve to ethanol and with 21-day IEE N per donor per replicate.** A spontaneously active neuron is defined as an ROI with at least one Ca^2+^ spike during the baseline acquisition period (3 minutes in ACSF). Refers to Figure 5D-G.

|  | | **Iso-CTL** | | **↑GIRK2** | |
| --- | --- | --- | --- | --- | --- |
| **Donor** | **Replicate** | **Naïve** | **21-Day IEE** | **Naïve** | **21-Day IEE** |
| 553 | 1 | 250 | 230 | 673 | 297 |
|  | 2 | 209 | 349 | 195 | 274 |
|  | 3 | 272 | -- | 183 | -- |
| 2607 | 1 | 118 | 315 | 197 | 48 |
|  | 2 | 64 | 58 | 31 | 96 |
|  | 3 | -- | -- | -- | 112 |
| 12455 | 1 | 266 | 184 | 51 | 98 |
|  | 2 | 232 | 83 | 178 | -- |
|  | 3 | 71 | -- | -- | -- |
| 9429 | 1 | 356 | 117 | 267 | 100 |
|  | 2 | 97 | 62 | 207 | -- |
| BJ | 1 | 205 | 357 | 147 | 86 |
|  | 2 | 283 | 102 | 85 | -- |
|  | 3 | -- | 126 | 70 | -- |
| Number of replicates/group (points plotted) | | 13 | 12 | 13 | 9 |
| TOTAL ROIs | | 1942 | 1404 | 1772 | 540 |
