## Extended Data Table 6-1 for "Upregulated GIRK2 counteracts ethanol-induced changes in excitability & respiration in human neurons"

**Extended Data Table 6-1: Total number of active neurons per donor, per replicate in iso-CTL and ↑GIRK2 cohorts naïve to ethanol and with 21-day IEE.** An active neuron is defined as an ROI with at least one Ca^2+^ spike during the duration of the entire experiment, across all epochs. Refers to Figure 6D.

|  | | **Iso-CTL** | | **↑GIRK2** | |
| --- | --- | --- | --- | --- | --- |
| **Donor** | **Replicate** | **Naïve** | **21-Day IEE** | **Naïve** | **21-Day IEE** |
| 553 | 1 | 299 | 230 | 691 | 297 |
|  | 2 | 272 | 349 | 183 | 274 |
|  | 3 | 209 | -- | 195 | -- |
| 2607 | 1 | 120 | 363 | 208 | 69 |
|  | 2 | 77 | 63 | 35 | 108 |
|  | 3 | -- | -- | -- | 126 |
| 12455 | 1 | 320 | 213 | 56 | 104 |
|  | 2 | 244 | 90 | 191 | -- |
|  | 3 | 77 | -- | -- | -- |
| 9429 | 1 | 378 | 162 | 270 | 139 |
|  | 2 | 100 | 64 | 226 | -- |
| BJ | 1 | 227 | 373 | 161 | 86 |
|  | 2 | 285 | 120 | 86 | -- |
|  | 3 | -- | 127 | 72 | -- |
| Number of replicates/group (points plotted) | | 13 | 12 | 13 | 9 |
| TOTAL ROIs | | 2608 | 2154 | 2374 | 1203 |
