## Extended Data Table 6-2 for "Upregulated GIRK2 counteracts ethanol-induced changes in excitability & respiration in human neurons"

**Extended Data Table 6-2: number of glutamate-active ROIs in iso-CTL and ↑GIRK2 cohorts naïve to ethanol and with 21-day IEE.** Refers to Figure 6E-H.

|  | | **Iso-CTL** | | **↑GIRK2** | |
| --- | --- | --- | --- | --- | --- |
| **Donor** | **Replicate** | **Naïve** | **21-Day IEE** | **Naïve** | **21-Day IEE** |
| 553 | 1 | 174 | 138 | 175 | 238 |
|  | 2 | 218 | 247 | 140 | 132 |
|  | 3 | 149 | -- | 301 | -- |
| 2607 | 1 | 44 | 145 | 110 | 19 |
|  | 2 | 60 | 25 | 22 | 32 |
|  | 3 | -- | -- | -- | 95 |
| 12455 | 1 | 133 | 86 | 27 | -- |
|  | 2 | 110 | 27 | 68 | 27 |
|  | 3 | 38 | -- | -- | -- |
| 9429 | 1 | 117 | 66 | 83 | 64 |
|  | 2 | 54 | 30 | 107 | -- |
| BJ | 1 | 150 | 272 | 93 | 40 |
|  | 2 | 136 | 56 | 50 | -- |
|  | 3 | -- | 32 | 43 | -- |
| Number of replicates/group (points plotted) | | 13 | 12 | 13 | 9 |
| TOTAL ROIs | | 1383 | 1124 | 1219 | 647 |
