## Extended Data Table 6-3 for "Upregulated GIRK2 counteracts ethanol-induced changes in excitability & respiration in human neurons"

**Extended Data Table 6-3: number of KCl- and glutamate-active neuronal ROIs in iso-CTL and ↑GIRK2 cohorts naïve to ethanol and with 21-day IEE.**

|  | | **Iso-CTL** | | **↑GIRK2** | |
| --- | --- | --- | --- | --- | --- |
| **Donor** | **Replicate** | **Naïve** | **21-Day IEE** | **Naïve** | **21-Day IEE** |
| 553 | 1 | 170 | 17 | 139 | 198 |
|  | 2 | 182 | 235 | 111 | 63 |
|  | 3 | 143 | -- | -- | -- |
| 2607 | 1 | 19 | 102 | 77 | 11 |
|  | 2 | 12 | 7 | 5 | 12 |
|  | 3 | -- | -- | -- | 23 |
| 12455 | 1 | 100 | 11 | 12 | 10 |
|  | 2 | 78 | 4 | 13 | -- |
|  | 3 | 4 | -- | -- | -- |
| 9429 | 1 | 47 | 27 | 11 | 42 |
|  | 2 | 8 | 1 | 16 | -- |
| BJ | 1 | 68 | 45 | 31 | 6 |
|  | 2 | 27 | 30 | 2 | -- |
|  | 3 | -- | 9 | 6 | -- |
| Number of replicates/group (points plotted) | | 13 | 12 | 13 | 9 |
| TOTAL ROIs | | 858 | 488 | 413 | 365 |
