## Extended Data Table 2-1 for "Upregulated GIRK2 counteracts ethanol-induced changes in excitability & respiration in human neurons"

| Pooled CRISPRa & Lenti | | |
| --- | --- | --- |
| Parameter | iso-CTL | ↑GIRK2 |
| N | 23 | 21 |
| Capacitance (pF) | 25.8 ± 2.6 | 23.3 ± 1.3 |
| Membrane Access Resistance (MΩ) | 13.7 ± 1.6 | 13.7 ± 1.3 |
| Rheobase (pA) | 42 ± 8 | 56 ± 10 |
| Action Potential Threshold (mV) | -29.8 ± 1.5 | -28.2 ± 1.5 |
| Resting Membrane Potential (mV) | -63.9 ± 1.6 | -67.4 ± 1.4 |
| CRISPRa (Donor 553-VPR) | | |
|  | iso-CTL  (Scramble gRNA) | ↑GIRK2  (*KCNJ6*gRNA) |
| N | 12 | 11 |
| Capacitance (pF) | 29.8 ± 3.4 | 23.7 ± 1.5 |
| Membrane Series Resistance (MΩ) | 15.4 ± 1.7 | 16.2 ± 1.7 |
| Rheobase (pA) | 35 ± 5 | 55 ± 13 |
| Action Potential Threshold (mV) | -31.1 ± 1.6 | -26.1 ± 2.4 |
| Resting Membrane Potential (mV) | -59.4 ± 2.0 | -69.4 ± 2.0 |
| Lenti (Donor 12455) | | |
|  | iso-CTL | ↑GIRK2  (*KCNJ6-*mCherry vector) |
| N | 11 | 10 |
| Capacitance (pF) | 17.9 ± 2.5 | 22.8 ± 2.2 |
| Membrane Series Resistance (MΩ) | 10.6 ± 3.0 | 10.7 ± 1.6 |
| Rheobase (pA) | 54 ± 18 | 56 ± 16 |
| Action Potential Threshold (mV) | -27.8 ± 2.9 | -29.4 ± 1.8 |
| Resting Membrane Potential (mV) | -63 ± 1.3 | -67.1 ± 2.6 |
