## Extended Data Table 3-1 for "Upregulated GIRK2 counteracts ethanol-induced changes in excitability & respiration in human neurons"

| **Gene name** | **log2(FC)** | **p-value** | **State** |  | **Gene name** | **log2(FC)** | **p-value** | **State** |
| --- | --- | --- | --- | --- | --- | --- | --- | --- |
| ABCB1 | 0.613229 | 0.008341 | Up |  | P2RY14 | 1.063311 | 0.029837 | Up |
| AC009014.3 | 0.62568 | 0.014469 | Up |  | PCK1 | 1.623995 | 8.70E-04 | Up |
| ACSBG1 | 0.894429 | 4.23E-05 | Up |  | PDK4 | 1.370501 | 1.45E-16 | Up |
| ACTBL2 | 1.230699 | 0.007814 | Up |  | PECAM1 | 0.803719 | 0.008728 | Up |
| ACVRL1 | 0.863414 | 0.004254 | Up |  | PHF11 | 0.674768 | 2.49E-05 | Up |
| ADGRF5 | 1.620721 | 2.49E-05 | Up |  | PIK3R6 | 1.006461 | 0.028872 | Up |
| ADPRH | 0.644971 | 0.002929 | Up |  | PLAT | 0.686664 | 1.70E-05 | Up |
| ADRA2B | 1.436806 | 3.63E-06 | Up |  | PLVAP | 1.297156 | 0.033623 | Up |
| AGT | 0.503381 | 0.001882 | Up |  | PPY | 0.755943 | 0.022059 | Up |
| AKR1C3 | 0.863601 | 1.30E-05 | Up |  | PRRX2 | 0.622337 | 0.03647 | Up |
| ALOX5 | 0.5138 | 0.043256 | Up |  | PTGDR2 | 0.599109 | 0.043526 | Up |
| ALOX5AP | 1.03469 | 0.021631 | Up |  | PTGER1 | 0.549397 | 0.007215 | Up |
| ALX4 | 1.264565 | 9.90E-06 | Up |  | PTGES | 1.173676 | 8.45E-08 | Up |
| ANGPTL4 | 4.056351 | 1.60E-29 | Up |  | PTPRB | 0.622809 | 0.038736 | Up |
| ANKH | 0.534896 | 1.11E-05 | Up |  | PYY | 1.04562 | 0.005331 | Up |
| ANXA13 | 0.890029 | 0.002197 | Up |  | RARRES1 | 1.163044 | 2.22E-04 | Up |
| AOC3 | 0.711753 | 0.003007 | Up |  | RASSF5 | 0.749568 | 1.83E-04 | Up |
| ARHGAP25 | 0.661258 | 0.026731 | Up |  | RGS5 | 0.565587 | 0.014736 | Up |
| ARHGDIB | 1.056393 | 7.78E-05 | Up |  | SELL | 0.712524 | 0.006767 | Up |
| ARSI | 0.6974 | 5.29E-05 | Up |  | SH3TC1 | 0.761453 | 5.94E-06 | Up |
| ATP8B1 | 0.555686 | 0.004738 | Up |  | SLC12A8 | 0.548772 | 7.99E-04 | Up |
| BATF2 | 0.738948 | 0.047386 | Up |  | SLC2A5 | 0.922765 | 7.78E-05 | Up |
| BDKRB1 | 0.789372 | 0.015793 | Up |  | SLC2A7 | 1.17412 | 0.006125 | Up |
| BHMT | 0.761408 | 0.014076 | Up |  | SMIM10 | 0.541046 | 0.033926 | Up |
| C10orf10 | 0.678214 | 7.49E-06 | Up |  | SMIM3 | 0.56691 | 1.76E-05 | Up |
| C1QTNF1 | 0.632175 | 5.58E-05 | Up |  | SMOC2 | 0.584815 | 0.003642 | Up |
| C3AR1 | 1.064852 | 0.006635 | Up |  | SPATA22 | 1.134736 | 0.019329 | Up |
| C7 | 0.979889 | 0.01552 | Up |  | SPP1 | 0.512751 | 0.01334 | Up |
| C7orf33 | 1.200112 | 0.001534 | Up |  | SPTLC3 | 0.64107 | 0.00698 | Up |
| CALCB | 0.839313 | 0.027652 | Up |  | SRPX2 | 0.563837 | 0.006617 | Up |
| CARD6 | 0.634391 | 0.046851 | Up |  | STEAP1 | 0.530143 | 0.018719 | Up |
| CCR1 | 0.869432 | 0.021356 | Up |  | STEAP4 | 1.196835 | 0.004299 | Up |
| CHI3L1 | 1.304242 | 2.16E-05 | Up |  | STOM | 0.585221 | 2.61E-04 | Up |
| CHRNA9 | 0.816685 | 1.17E-05 | Up |  | TAPBPL | 0.516724 | 0.006503 | Up |
| CLEC2B | 0.781784 | 0.006767 | Up |  | TEX14 | 0.534242 | 0.014396 | Up |
| CLUL1 | 0.869053 | 0.001814 | Up |  | TGM2 | 0.968232 | 8.45E-08 | Up |
| CMKLR1 | 0.70412 | 0.004516 | Up |  | TIMP1 | 0.713865 | 0.003334 | Up |
| CPA1 | 0.824748 | 0.002471 | Up |  | TKTL1 | 0.786881 | 0.001442 | Up |
| CPED1 | 0.54152 | 0.009128 | Up |  | TLR7 | 1.094965 | 0.010266 | Up |
| CPT1A | 0.85834 | 3.44E-11 | Up |  | TMEM144 | 0.646811 | 4.72E-05 | Up |
| CRABP2 | 0.547229 | 0.034018 | Up |  | TMEM204 | 0.714975 | 0.014736 | Up |
| CRISPLD2 | 0.670963 | 0.002368 | Up |  | TNFRSF8 | 1.092917 | 2.49E-05 | Up |
| CRYGN | 0.92461 | 0.003886 | Up |  | TSKS | 1.163065 | 0.002615 | Up |
| CTCFL | 1.294727 | 1.37E-04 | Up |  | ABCG1 | -0.7347 | 2.37E-04 | Down |
| CTHRC1 | 0.759595 | 6.16E-04 | Up |  | ABI3BP | -0.71264 | 2.47E-04 | Down |
| CTSZ | 0.577351 | 0.002468 | Up |  | ADM2 | -0.64435 | 8.04E-04 | Down |
| CXCL12 | 0.56748 | 0.019244 | Up |  | ADRA1B | -0.88833 | 0.042658 | Down |
| CYP26A1 | 1.382086 | 2.88E-06 | Up |  | AHNAK2 | -0.58059 | 0.006209 | Down |
| DACT2 | 0.690376 | 7.67E-04 | Up |  | ARHGAP9 | -0.65135 | 0.003962 | Down |
| DCSTAMP | 1.656884 | 6.01E-04 | Up |  | ASCL5 | -0.83835 | 0.034084 | Down |
| DHRS3 | 0.62705 | 3.39E-04 | Up |  | BMP7 | -0.50801 | 0.009128 | Down |
| DOK2 | 0.936548 | 0.034962 | Up |  | C1orf95 | -0.58339 | 0.003962 | Down |
| DPYS | 1.060336 | 5.59E-04 | Up |  | CBLN4 | -0.52207 | 0.00813 | Down |
| DUSP2 | 1.063815 | 5.08E-04 | Up |  | CLEC19A | -1.10846 | 0.010815 | Down |
| EGFLAM | 0.78556 | 0.001528 | Up |  | CNTNAP5 | -0.86035 | 0.025418 | Down |
| EMCN | 0.975288 | 0.001459 | Up |  | CRHR1 | -0.74295 | 0.048312 | Down |
| ENPEP | 0.531788 | 0.019448 | Up |  | CRYM | -0.58033 | 0.004804 | Down |
| ETV4 | 0.532009 | 0.003428 | Up |  | CST7 | -0.89258 | 0.049096 | Down |
| EVC2 | 0.567302 | 0.027176 | Up |  | DCC | -0.57556 | 0.00336 | Down |
| EXOC3L2 | 1.124024 | 8.45E-08 | Up |  | DPP6 | -0.63269 | 0.014721 | Down |
| FABP4 | 1.202082 | 1.58E-04 | Up |  | EPB41L4A | -0.50737 | 0.009815 | Down |
| FAM20C | 0.514158 | 0.002536 | Up |  | EPHA10 | -0.5206 | 0.040493 | Down |
| FES | 0.576556 | 6.54E-05 | Up |  | EPHB1 | -0.60961 | 2.93E-05 | Down |
| FGFBP2 | 0.553075 | 0.016929 | Up |  | FAM129A | -0.63377 | 0.001339 | Down |
| FMO2 | 0.603245 | 0.045512 | Up |  | FAM46C | -0.80505 | 0.002197 | Down |
| FMO3 | 0.875523 | 0.001165 | Up |  | FBN3 | -0.52159 | 7.58E-04 | Down |
| FMOD | 0.647739 | 0.027143 | Up |  | GCGR | -1.10712 | 0.029543 | Down |
| GABRR1 | 1.275394 | 8.45E-08 | Up |  | GPR1 | -0.68629 | 5.55E-04 | Down |
| GALNT5 | 0.613182 | 0.001094 | Up |  | GPR146 | -1.0386 | 0.019188 | Down |
| GDF3 | 0.602473 | 0.021795 | Up |  | GPR26 | -0.89042 | 0.014487 | Down |
| GGT5 | 0.518324 | 0.007624 | Up |  | GREM1 | -0.51443 | 0.00426 | Down |
| GLIPR1 | 0.673103 | 1.82E-05 | Up |  | GRIN2B | -0.557 | 0.006125 | Down |
| GNA14 | 0.607291 | 0.035319 | Up |  | GRM4 | -0.5235 | 0.033808 | Down |
| GPR81 | 0.823865 | 0.048305 | Up |  | HES2 | -0.57391 | 4.42E-04 | Down |
| HOPX | 0.643527 | 7.60E-05 | Up |  | IL11 | -0.52926 | 0.02141 | Down |
| HOXB1 | 1.048777 | 9.55E-05 | Up |  | INHBE | -0.78838 | 0.022485 | Down |
| HP | 1.494634 | 7.42E-06 | Up |  | IQUB | -0.5017 | 0.008891 | Down |
| HRASLS5 | 0.556988 | 1.05E-04 | Up |  | ITIH5 | -0.59288 | 5.02E-04 | Down |
| HSPA6 | 0.830642 | 0.029837 | Up |  | KCNH1 | -0.58452 | 0.029651 | Down |
| IER3 | 0.526771 | 1.05E-04 | Up |  | KCNJ9 | -0.58034 | 0.012498 | Down |
| IFITM1 | 0.589311 | 0.003514 | Up |  | KCNQ3 | -0.52507 | 0.002197 | Down |
| IFITM2 | 0.67447 | 7.78E-05 | Up |  | KLF4 | -0.62996 | 0.016486 | Down |
| IFITM3 | 0.551147 | 5.68E-04 | Up |  | KLHDC7B | -1.59156 | 7.68E-05 | Down |
| IGFBP6 | 0.510583 | 0.01634 | Up |  | LA16c-380H5.3 | -0.94809 | 0.007514 | Down |
| IL1R1 | 0.738508 | 3.58E-04 | Up |  | LINGO3 | -0.69949 | 0.017779 | Down |
| IL1R2 | 1.210024 | 3.46E-04 | Up |  | NANOG | -0.89696 | 0.029837 | Down |
| IL7R | 0.911012 | 0.001534 | Up |  | NEURL3 | -0.70537 | 0.048037 | Down |
| INMT | 0.71909 | 0.001791 | Up |  | NFATC2 | -0.60116 | 0.014537 | Down |
| ITK | 0.994863 | 0.01472 | Up |  | NGF | -0.67456 | 0.003735 | Down |
| JAK3 | 0.580019 | 0.00319 | Up |  | NOG | -0.66182 | 0.006635 | Down |
| KCNE4 | 0.640271 | 7.78E-05 | Up |  | NPAS4 | -0.68027 | 0.041128 | Down |
| KLF10 | 0.528958 | 3.99E-06 | Up |  | NTRK2 | -0.60675 | 0.011189 | Down |
| KLF9 | 0.574185 | 4.58E-05 | Up |  | OSCAR | -0.88401 | 4.21E-04 | Down |
| KLHL6 | 1.244916 | 8.73E-04 | Up |  | PDE1C | -0.51898 | 0.003447 | Down |
| LAPTM5 | 0.642622 | 0.041128 | Up |  | PDE6C | -0.52256 | 0.047488 | Down |
| LBP | 1.045754 | 0.028031 | Up |  | PIK3CD | -0.50285 | 1.73E-04 | Down |
| LCP2 | 0.606477 | 0.033999 | Up |  | PLA2G3 | -0.67896 | 0.022417 | Down |
| LGALS12 | 1.035902 | 4.04E-05 | Up |  | POTEF | -0.58597 | 0.045944 | Down |
| LGALS9 | 0.862623 | 9.90E-06 | Up |  | PPL | -0.6046 | 2.61E-04 | Down |
| LRG1 | 1.234794 | 0.002978 | Up |  | PPP1R1B | -0.53202 | 0.026854 | Down |
| LRRC15 | 0.681639 | 0.01017 | Up |  | PRELP | -0.61993 | 0.019746 | Down |
| LRRC32 | 0.900335 | 3.63E-06 | Up |  | PTPRT | -1.40333 | 0.012116 | Down |
| LSP1 | 0.623594 | 0.002633 | Up |  | RBFOX1 | -0.67978 | 0.009672 | Down |
| LTBR | 0.823485 | 5.08E-04 | Up |  | RP4-614O4.11 | -0.97537 | 0.040118 | Down |
| LY96 | 0.842502 | 0.003781 | Up |  | SCARA5 | -0.78556 | 0.014076 | Down |
| M1AP | 0.765764 | 0.004972 | Up |  | SLC13A5 | -1.18499 | 0.002076 | Down |
| MEDAG | 0.942956 | 2.92E-04 | Up |  | SREBF1 | -0.51579 | 2.01E-06 | Down |
| MGP | 0.938778 | 4.01E-06 | Up |  | SSPO | -0.78087 | 0.00336 | Down |
| MMP13 | 1.904175 | 2.88E-06 | Up |  | SST | -0.6602 | 0.048483 | Down |
| MMP9 | 0.763591 | 0.024643 | Up |  | STAC2 | -0.53937 | 0.011994 | Down |
| MPO | 0.619998 | 0.029998 | Up |  | STC2 | -0.70909 | 0.00336 | Down |
| NINJ2 | 0.938053 | 0.027012 | Up |  | TH | -0.62768 | 0.014514 | Down |
| NNMT | 0.646371 | 0.005634 | Up |  | TMPRSS7 | -0.67393 | 0.031661 | Down |
| OAS1 | 0.687775 | 0.042597 | Up |  | TNR | -0.64601 | 0.014801 | Down |
| OAS2 | 0.770632 | 0.01162 | Up |  | TRIB3 | -0.80696 | 9.90E-06 | Down |
| OASL | 0.826176 | 0.033693 | Up |  | TTC34 | -0.68798 | 0.019269 | Down |
| OTOGL | 0.637818 | 0.033749 | Up |  | VRTN | -0.97364 | 0.004254 | Down |
